# Circadian diversity in Swedish *Arabidopsis* accessions is associated with naturally occurring genetic variation in *COR28*

**DOI:** 10.1101/665455

**Authors:** Hannah Rees, Ryan Joynson, James K.M. Brown, Anthony Hall

## Abstract

Circadian clocks have evolved to resonate with external day and night cycles. However, these entrainment signals are not consistent everywhere and vary with latitude, climate and seasonality. This leads to divergent selection for clocks which are locally adapted. To investigate the genetic basis for this, we used a Delayed Fluorescence (DF) imaging assay to screen 191 naturally occurring Swedish *Arabidopsis* accessions for their circadian phenotypes. We demonstrate period variation with both latitude and sub-population. Several candidate loci linked to period, phase and Relative Amplitude Error (RAE) were revealed by genome-wide association mapping and candidate genes were investigated using TDNA mutants. We show that natural variation in a single non-synonymous substitution within *COR28* is associated with a long-period and late-flowering phenotype similar to that seen in TDNA knock-out mutants. *COR28* is a known coordinator of flowering time, freezing tolerance and the circadian clock; all of which may form selective pressure gradients across Sweden. Finally, we tested circadian variation under reduced temperatures and show that fast and slow period phenotypic tails remain diverged and follow a distinctive ‘arrow-shaped’ trend indicative of selection for a cold-biased temperature compensation response.

## Introduction

Plants are highly adapted to survive and exploit the daily fluctuations in light and temperature experienced as the earth spins on its axis. The circadian clock plays an intrinsic role in this; integrating temporal cues from the environment to inform photosynthetic, metabolic and developmental processes(1,2). A robustly oscillating circadian clock which is highly synchronised to external day-length contributes to the overall fitness of the plant, giving it an edge over competitors, predators and pathogens(3–6).

*Arabidopsis thaliana* is a model plant system which has been extensively studied in circadian biology. The *Arabidopsis* clock is comprised of a series of interlocking negative transcriptional feedback loops connected by key activators that control the oscillation of clock gene expression(7). Key genes within this network include *CIRCADIAN CLOCK ASSOCIATED 1 (CCA1)* and *LATE ELONGATED HYPOCOTYL (LHY)* which transcriptionally repress *TIMING OF CAB EXPRESSION (TOC1)*(8). Downstream genes interact with the clock to communicate circadian rhythmicity to physiological outputs. Examples include photoperiodic regulation of flowering time and the temporal gating of cold acclimation responses(9). Flowering under long days is instigated through accumulation of the floral promoter CONSTANS (CO) controlled by the circadian clock component GIGANTEA(10). In winter-annuals, flowering is also dependent on the vernalization response of *FLOWERING LOCUS C (FLC)* and *FRIGIDA (FRI)*(11,12). Cold tolerance is diurnally activated through alternative splicing of *CCA1* which co-regulates the expression of *COLD REGULATED (COR)* genes alongside light and temperature stress pathways(9,13). Circadian rhythms can be quantified by their period (the time taken to complete one cycle), their phase (the time of day in which the rhythm peaks), their amplitude (the change in intensity from a baseline) and their Relative Amplitude Error (RAE) which is the amplitude error divided by the overall amplitude of the rhythm and can be equated to rhythm robustness.

The endogenous core circadian network is entrained by external stimuli; most notably light and temperature. Day-length, light composition and light intensity have all been shown to affect circadian rhythms(14–17). Temperature also has a well-documented effect on entrainment of circadian rhythms(18–20) with a degree of period shortening expected under higher temperatures. Circadian rhythms are said to be ‘temperature compensated’; they resist large changes in period-length in response to temperature(21). The extent of temperature compensation has been shown to vary between accessions(20,22). Rhythm robustness is also strongly affected by temperature, although the temperature which produces the most rhythmic oscillations appears to be dependent on the species and circadian assay used (22,23). Plants with clocks which resonate with environmental conditions are typically fittest, however it has been suggested that in climates with large seasonality there may be a compromise for clocks which are adaptable to rapidly changing day-lengths(6).

Natural populations of several important model organisms that exhibit extensive diversity in their circadian behavior have been documented. Pupal eclosion rhythms of Drosophila are latitude dependent; with shorter rhythms, earlier phases and less robust rhythms observed in Northern latitudes(24,25). In *Arabidopsis*, Michel et al. conducted a global study of leaf movement rhythms in 150 accessions and found that day-length of origin country correlated positively with period, but not phase or amplitude. They also identified several loci in the *TOC1/PRR* family which determined natural variation in period, phase and amplitude independently(6). Other investigations have shown allelic diversity of several clock related genes including *FLC*(26) *GI*(27) and *EARLY FLOWERING 3 (ELF3)*(28) which contribute to natural circadian phenotypes without fully disrupting the clock mechanism. The positive relationship with period and latitude has been replicated in tomato(29), soybean and annual populations of *Mimulus guttatus*(30) and gives a strong indication that this is an adaptive phenotype.

In this study, we have focused on a collection of *Arabidopsis* accessions from across a large latitudinal range in Sweden; a country with variations in climatic, anthropogenic and day-length factors all potentially influencing clock adaption. Northern latitudes around 63°N have permanent snow cover during winter months and the growing season is much cooler and shorter than in the South. At the solstice, there is almost 3 hours difference in day-length between the North and South. This divergence of climate has led to selection for ecotypes with adapted growth and flowering strategies(31,32). Analysis of the global population structure of *Arabidopsis* accessions has previously identified Swedish accessions as being genetically distinct from the wider population, with further differentiation within the country between North and South(33–36). Accessions from South Sweden have high genetic diversity within a relatively small area, perhaps suggesting a historic emigration from central Europe following glacial retreat(37). Northern Swedish accessions have lower genetic diversity but larger genome sizes(35) and also carry a surprising reservoir of drought tolerance genes(38). Completion of the 1001 genomes project in 2006 has facilitated a recent expansion in *Arabidopsis* genome wide association (GWA) studies made possible through the provision of high-quality re-sequenced genotype data. These accessions are publicly available, geo-referenced and genetically inbred making it easy for researchers to perform experiments over several generations under a variety of conditions(39). Many of the accessions in this study have also been characterized for other phenotypic traits such as seed dormancy(40), flowering time(41), and freezing tolerance(42).

Here we use delayed fluorescence (DF) imaging and association mapping to reveal natural allelic diversity in several genes which contribute to circadian phenotypes. Delayed fluorescence levels are circadian regulated and reflect the changes in the photosynthetic state of photosystem II(43). DF imaging has been validated as a reliable and flexible tool to measure circadian rhythms in a range of plant models but has not previously been used for phenotyping of a large population of individuals on the scale required for genome-wide association mapping.

## Results

### Quantitative circadian variation across 191 Swedish *Arabidopsis* accessions

We used delayed fluorescence imaging to characterize circadian rhythms in groups of 14 day old seedlings entrained in 12h L 12h D cycles at 22°C and assayed under free-running conditions of constant light (L:L) at 22°C. Accession means and standard errors (SE) adjusted for experimental effects were obtained using linear mixed models for period, amplitude and RAE and a circular regression model for phase (see Extended Methods). We found a 4.42h difference between the mean period of the fastest (21.28h, SE=0.329) and slowest (25.70h, SE=0.355) accessions tested (Figure 1A). The mean phase of peak DF intensity for each accession occurred over the dark half of the cycle (12-24/0h) with a huge difference (10.32h) from the earliest peaking (12.21h, SE=0.58) to the latest (22.54h, SE=0.20) (Figure 1B). This variation in period and phase is consistent with data previously reported from a global collection of *Arabidopsis* accessions(6). We also measured the robustness of these rhythms by looking at their RAE values. The most rhythmic (lowest RAE) mean was 0.31 (SE=0.023) and the least rhythmic was 0.44 (SE=0.024) representing approximately 20% of the possible range of this trait in this study (Figure 1C). Accession means for circadian phenotypes can be viewed in Supplementary File 1. To test the effect of accession ID on each trait we performed a likelihood ratio test against a model which omitted accession ID as a variance component. Including accession ID in the mixed model had highly significant effects on period (χ^2^ (1 df)=455.57, *p<*0.0001) and RAE (χ^2^ (1)= 65.97, *p<*0.0001). Including Accession ID in the circular regression model also had a strongly significant effect on phase (χ^2^ (191)=407, *p<*0.0001). Amplitude was analysed as a log_10_ transformation (see Extended Methods) and there was no significant effect of accession ID on Log10Amplitude (χ^2^ (1)= 0.19, *p=*0.6). With this in mind, we dropped Amplitude as a trait of interest for the temperature and mutant screening experiments. REML output tables (Supplementary Tables 1-3), model checking graphs (Supplementary Figures 1-3) and log-likelihood results (Supplementary Tables 4-7) are available to view.

**Figure 1.**
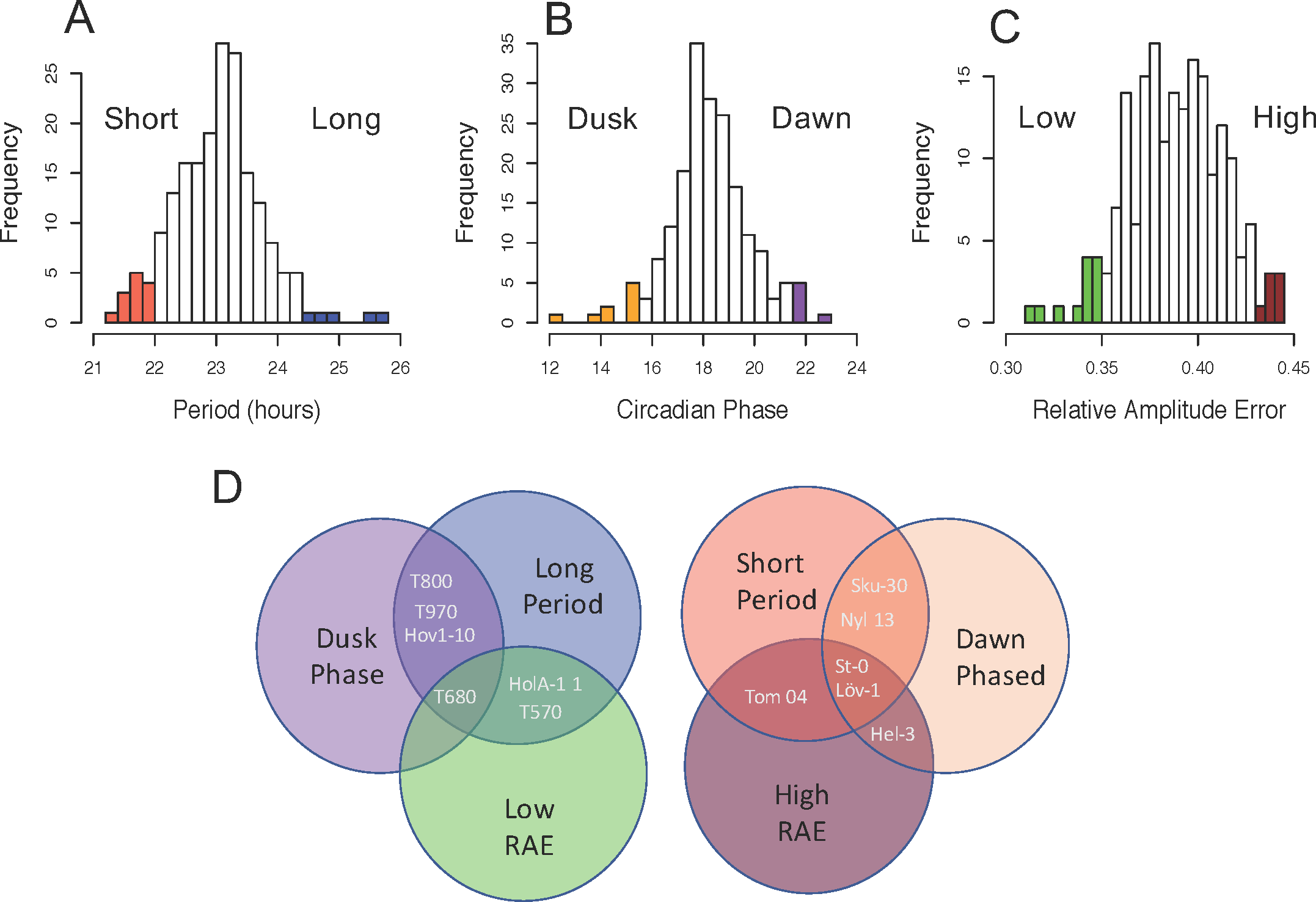
Circadian variation in 191 naturally occurring accessions from Sweden. Delayed fluorescence rhythms were characterised by period, phase and RAE and show significant variation (A-C). Colours represent the 10 most extreme accessions for each phenotype which we use as tail groups for further analysis. Some accessions belong to multiple tail groups as is shown in D. This reflects the strong correlation between circadian characteristics (See Supplementary Figure 4). Number of individual wells contributing to the mean of each accession ranged from 4 to 18, with each well representing the mean rhythms of approximately 15 seedlings.

**Figure 2.**
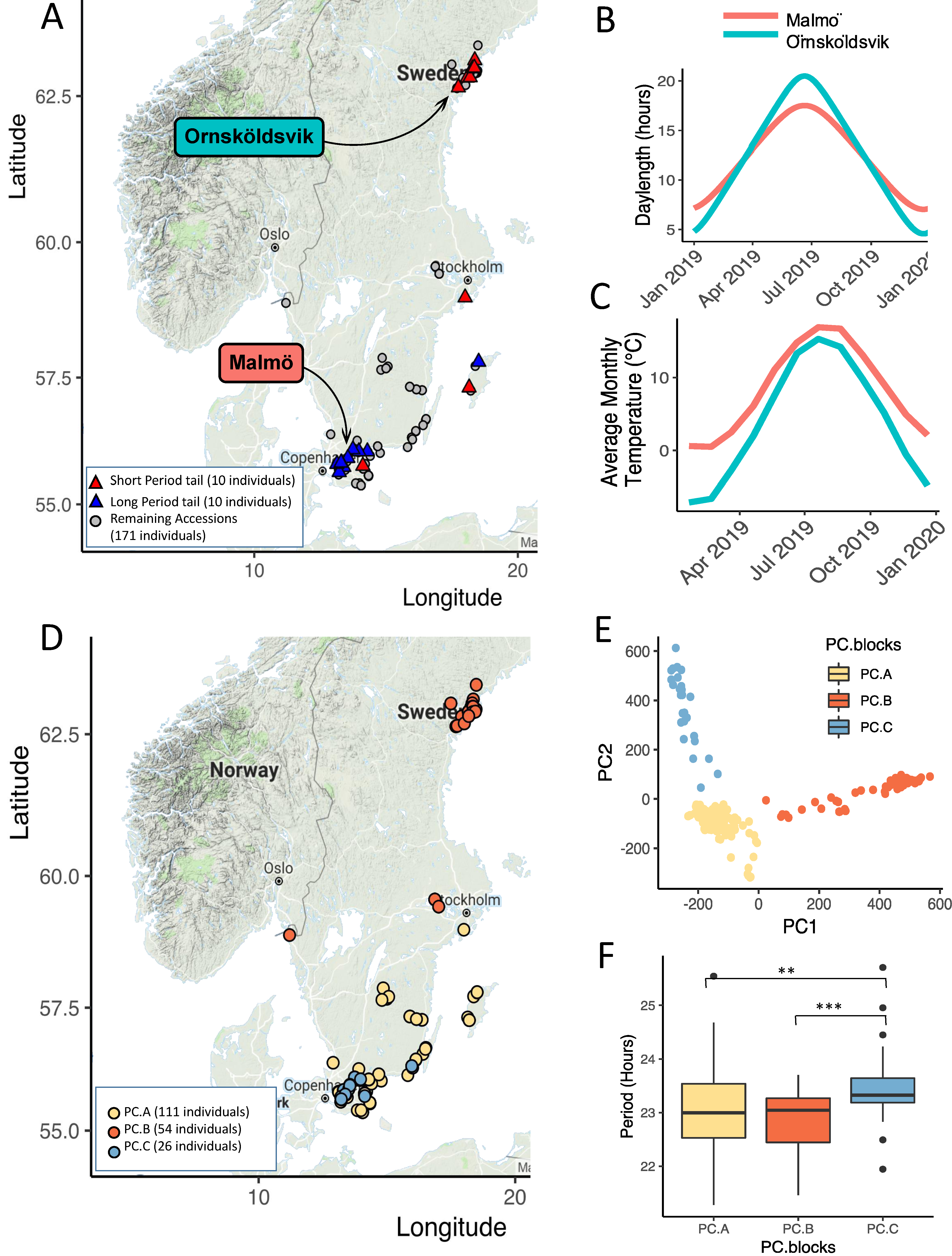
Period phenotypes are linked by both the geography and the genetic relatedness of accessions across Sweden. Figure A shows the location of origin of the 10 accessions with the shortest (red triangles) and longest (blue triangles) periods. The longest periods are found near the city of Malmo where day length and temperature fluctuates less throughout the year than in Ornskoldsvik where periods tend to be shorter (B,C). Day-length and temperature averages were downloaded from timeanddate.com and are based on predictions for 2019 (60). PC analysis revealed a sub-clade within the Southern Swedish accessions; PC.C coloured light blue in plots D-F. This was used to distinguish accessions with significantly longer period phenotype than in the other PC groups; PC.A which represents the remaining southern accessions in yellow and PC.B which represents the northern accessions in red. Map figures were created using the ggmaps package in R using Google maps (accessed 2018)(56).

We observed a strong correlation between the period, RAE and phase of each accession (Supplementary Figure 4), especially between period and phase (Adjusted R^2^ =0.43, *p<*0.0001). We are unaware of any previously published correlation between natural variation in period and RAE. In this study, we found a highly significant negative relationship with longer periods having lower RAE scores (Adjusted R^2^ =0.1, *p<*0.0001). We chose 10 accessions from the tails of the distributions of mean period, phase and RAE to create “phenotypic tail” groups representing the extremes of each trait. Some accessions represented two or more tail groups and could be split into two master groups of: long period, dusk phased, low RAE accessions and short period, dawn phased, high RAE accessions as explained in Figure 1D. These phenotypic tail groups were used for the temperature experiments described later in this paper (accessions in each tail are listed in Supplementary Tables 8-10).

Our next question was whether other quantified traits co-varied with our circadian data. We used several previously published datasets to examine possible phenotypic relationships between our traits and flowering time(41,44), seed dormancy(40) and freezing tolerance(42) (Supplementary Table 11). Only flowering time data from Li et al. 2010 showed any significant correlation with period and phase (*p<*0.01) based on flowering times recorded in conditions mimicking natural Swedish and Spanish climates(44). However no significant correlations were found between flowering times recorded under long-days under either 10°C or 16°C by Horton et.al 2016, indicating that the relationship is highly dependent on the exact experimental conditions used.

We then investigated whether circadian traits varied according to the original location of the accessions. Accessions with the shortest periods tended to be found in Northern and mid-latitude regions and the longest periods were found in the south (Figure 2A). Period was found to be significantly correlated with longitude and latitude (*p<*0.001), however a considerable amount of variation remained unexplained (Adjusted R^2^ <0.1) (Supplementary Figure 5). Following on from this, we asked the question whether population sub-structure could better explain the distribution of period phenotypes. We re-classified the population into three groups split across the axes of the first two principal components (which together explain over 15% of the total genetic variation) (Figure 2D and E). We found that period is significantly longer in group ‘PC.C’ (Figure 2F) (F(2,188)=8.39, TukeyHSD for PC.A and PC.C *p<*0.01, One-way ANOVA). PC.C represents a cluster of accessions from the South of Sweden which are genetically distinct from other accessions found in the same geographic region. PC.A largely consisted of the remaining southern accessions from this study which had mean periods similar to those obtained from Northern accessions (PC.B). PC.C accessions also have significantly earlier phase peaks than those in PC.A (One-way ANOVA, F(2,188)=6.9149, TukeyHSD for PC.A and PC.C *p<*0.001).

### GWA mapping reveals loci associated with natural circadian variation

We used the online GWA-portal to perform association mapping on the three circadian traits found to be significantly affected by genetic variation.(45) We used an Accelerated Mixed Model (AMM) to account for population structure, although the other association models (linear and non-parametric) gave similar results (See Supplementary Figure 6). Pseudo-heritability estimates for each trait were given as an output of the GWA analysis: period=71%, RAE=37% and phase=13%.

GWA identified multiple genomic regions associated with period, phase and RAE as shown in Figure 3. We used –log10(*p*) 6.5 as an arbitrary p-score cut-off to select SNPs for further investigation. This threshold is conservative compared to several other previously published studies in *Arabidopsis*(46–48). We investigated known core circadian and flowering time genes to see whether these were significantly associated with any of the traits measured. Other than *ELF3* (see below), none of these genes fell within the window of association for significant SNPs. Genes in regions 30kb upstream or downstream of the most significant SNPs were considered as potential gene candidates and were selected for further analysis based on their previously attributed functionality and designated GO term. Genes involved in circadian rhythms, flowering time or chloroplast regulation were given prevalence (details in Supplementary File 2). The most significant associations had three SNPs with a –log10(*p*) score of 10.4-11.5, found on chromosome 4 associated with period variation. Within this interval we identified a non-synonymous SNP in the gene *COLD-REGULATED GENE 28*, a gene that has previously been identified as a negative regulator of several core clock genes (*PRR7, TOC1, PRR5* and *ELF4*) and is also implicated in the trade-off between flowering-time and freezing tolerance(49,50). The substitution resulted in a tryptophan (W) to serine (S) amino acid change at position 58 within the second exon of *COR28* (Figure 4A). This had a SIFT score of 0 indicating a highly probable deleterious effect on protein function. 16 accessions in this study had the minor allele, all found in the South of Sweden. The 58S accessions had a period 1.29h longer than the 58W accessions (t(17.4)=-7.46, *p<*0.001, Welch Two Sample t-test) (see Figure 4B) and mostly belonged to the genomic sub-group PC.A. The long-period of cluster PC.C therefore cannot be explained by variation in this SNP. We used the online tool Polymorph 1001 to look for other variants with the serine substitution and found only 5 other variants not assayed in this study (making 21 in total), all of which were also from the South of Sweden (Figure 4C).

**Figure 3.**
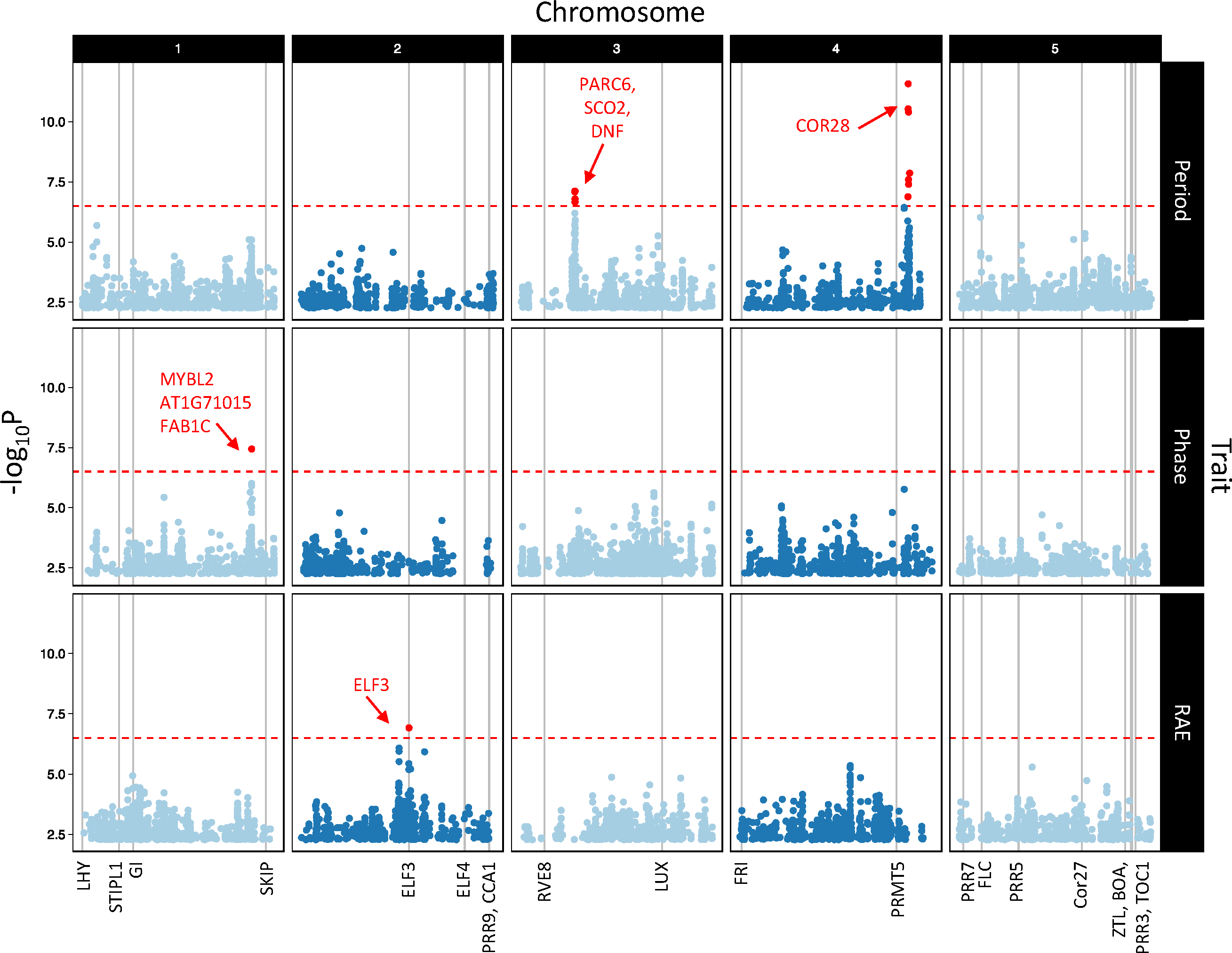
GWA analysis reveals candidate genes underlying natural variation in DF circadian phenotypes. Manhattan plots show AMM derived SNP associations with period (top panel), phase (middle panel) and RAE (bottom panel). Data shown has >5% MAF and reflects the 10,000 highest – log10(p) values. Gene candidates selected for further analysis are labelled in red with arrows. The red dotted line indicates the 6.5 –log10(p) arbitrary threshold used for this study. Vertical grey lines are labelled under the x axis and represent positions of known circadian and flowering time genes with altered circadian phenotypes previously reported in mutants.

**Figure 4.**
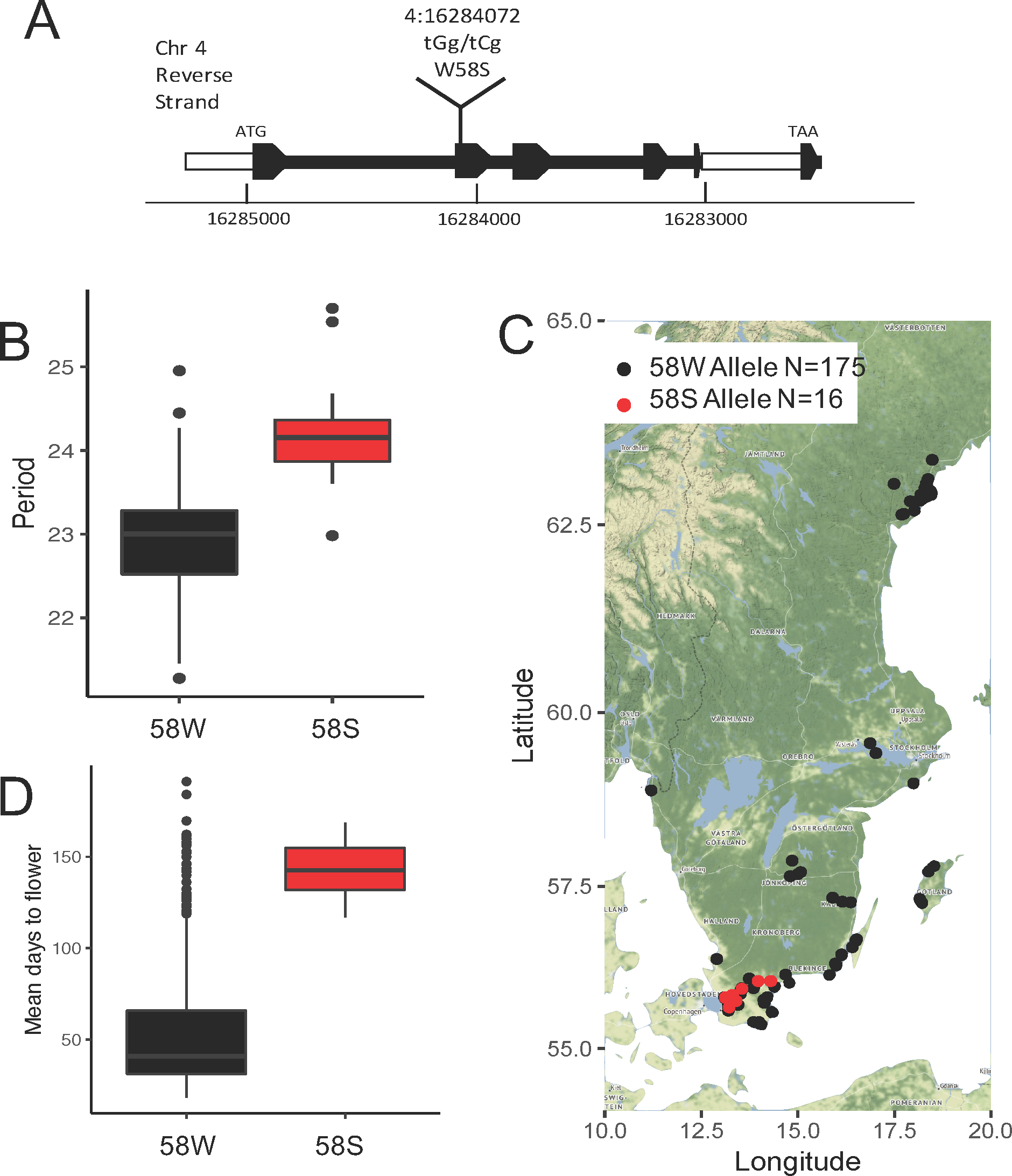
Natural Variation in COR28 is associated with covariation in period and flowering time. The SNP in COR28 associated with period variation causes a tryptophan (W) to serine (S) amino acid change within the second exon (A). Accessions carrying the 58S allele had periods 1.29h longer on average than those with the 58W allele based on the 191-accession data from this study (B). Out of the 191 accessions originally phenotyped, 16 had the 58S allele (globally, 5 further accessions were identified) and all were found in the south of Sweden (C). Accessions carrying the 58S allele also had flowering times delayed by 89.23 days compared to those carrying the 58W allele (data shown is based on flowering times from plants grown under Swedish conditions previously published in Li et al. 2010 (D).

We used flowering time data by Li et al. 2010(44) and Sasaki et al. 2015(41) to look for flowering time differences between the accessions carrying the *COR28*-58W major allele and the -58S minor allele (see Table 1). We found that the accessions with *COR28*-58S had significantly extended flowering times under simulated seasons for Sweden and Spain (Li) and under a 10°C and 16°C long-day temperature regime (Sasaki et al. 2016) (see Figure 4D). This complements previous findings that *COR28* is a flowering promoter and that modifications to this gene increase flowering time as well as lengthening period.

**Table 1.**
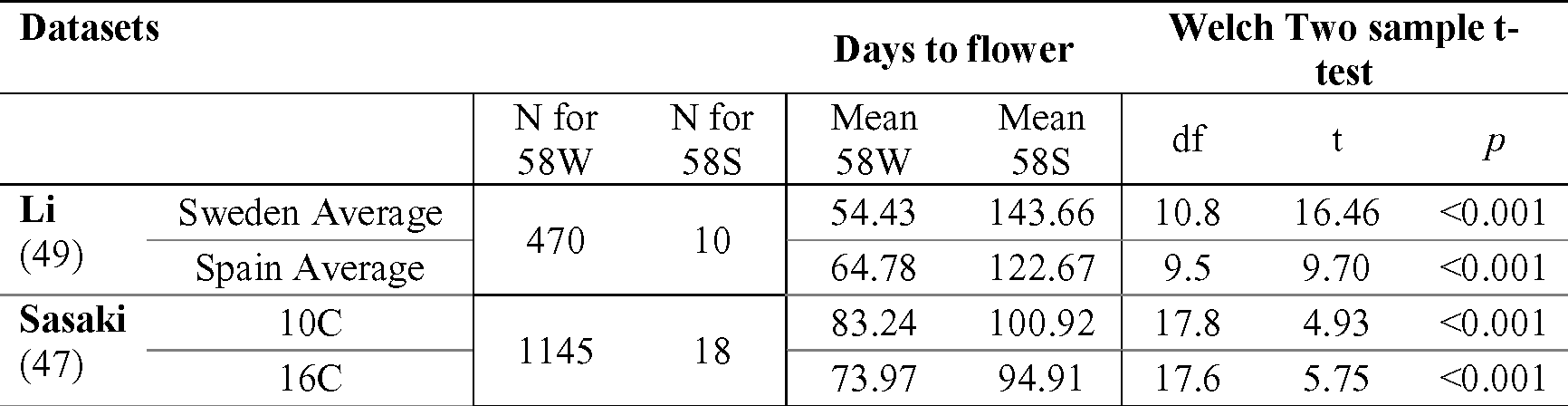
Longer flowering time correlation with COR28 W58S Allele.

N=Number of accessions used for each COR28 allele. Flowering data means reflect an average of all replicates within each study by Li et al. 2010 or Sasaki et al. 2016.

Other significant SNPs associated with period (chromosome 3) and phase (chromosome 1) identified gene candidates involved in chloroplast function (*PARC6, SCO2, DNF, AT1G71015* and *FAB1C*). The DF circadian output we used to assay these traits is based on oscillating activation phases of PSII and therefore natural variation in genes regulating chloroplast functionally could also affect the DF output in these candidates. A RAE associated SNP on chromosome 2 was found 5726bp upstream from a SNP in ELF3 previously characterized as the ELF3-Sha allele. Our associated SNP was found to be under strong linkage disequilibrium with the ELF3-Sha allele (R^2^=0.86). The alanine-to-valine transition in amino acid position 362 has been associated with naturally occurring alterations to periodicity and robustness in accessions from Central Asia, specifically Tajikistan(28). Here, 13 Swedish accessions were shown to carry the ELF-Sha allele and had mean RAE ratios 0.032 higher on average than for the other accessions. We substantiate evidence that ELF3-Sha accessions have lower rhythmicity and extend the global range of this allele into Northern Sweden. A surprising kinship between *Arabidopsis* accessions from Northern Sweden and Central Asia has been previously demonstrated through analysis of global population structure and indicates that the presence of this allele has not evolved convergently between the two populations(33).

### Validation of candidate genes

11 mutant lines representing eight gene candidates were genotyped to confirm homozygous mutations before being bulked for seed. Confirmed mutants were then assayed for their circadian rhythms using DF under the same conditions as for the 191 accession screen. Mutant details and genotyping results can be viewed in Supplementary File 2. *Cor27* mutants and the double mutant *cor27-1/28-2* were also assayed as *cor27* is known to be partially redundant with *cor28*. Analysis of the DF rhythms confirmed that periods in *cor28* mutants (*cor28-1*=25.3h, SD=1.73; *cor28-2*=25.3h, SD=1.78) were significantly longer than their respective controls (23.9h, SD=1.62 and 24.0h SD=2.02) (Welch Two Sample t-test *p<*0.05) as shown in Table 2. This confirms results previously reported using leaf movement, qPCR and ProCCR2:LUC bioluminescence rhythms(49,50). *Cor27* mutants showed no significant difference in period length, however the double knock-out *cor27-1/28-2* had an exaggerated long period phenotype (26.4h, SD=2.37). The peak phases of *cor27, cor28* and especially *cor27-1/28-2* were also earlier than for Col-0, with *cor27-1/28-2* having a mean phase peak 2.6h earlier than the control. Other mutants characterized were not found to have significantly altered periods or phases compared to their WT controls (Supplementary Tables 12-14).

**Table 2.**
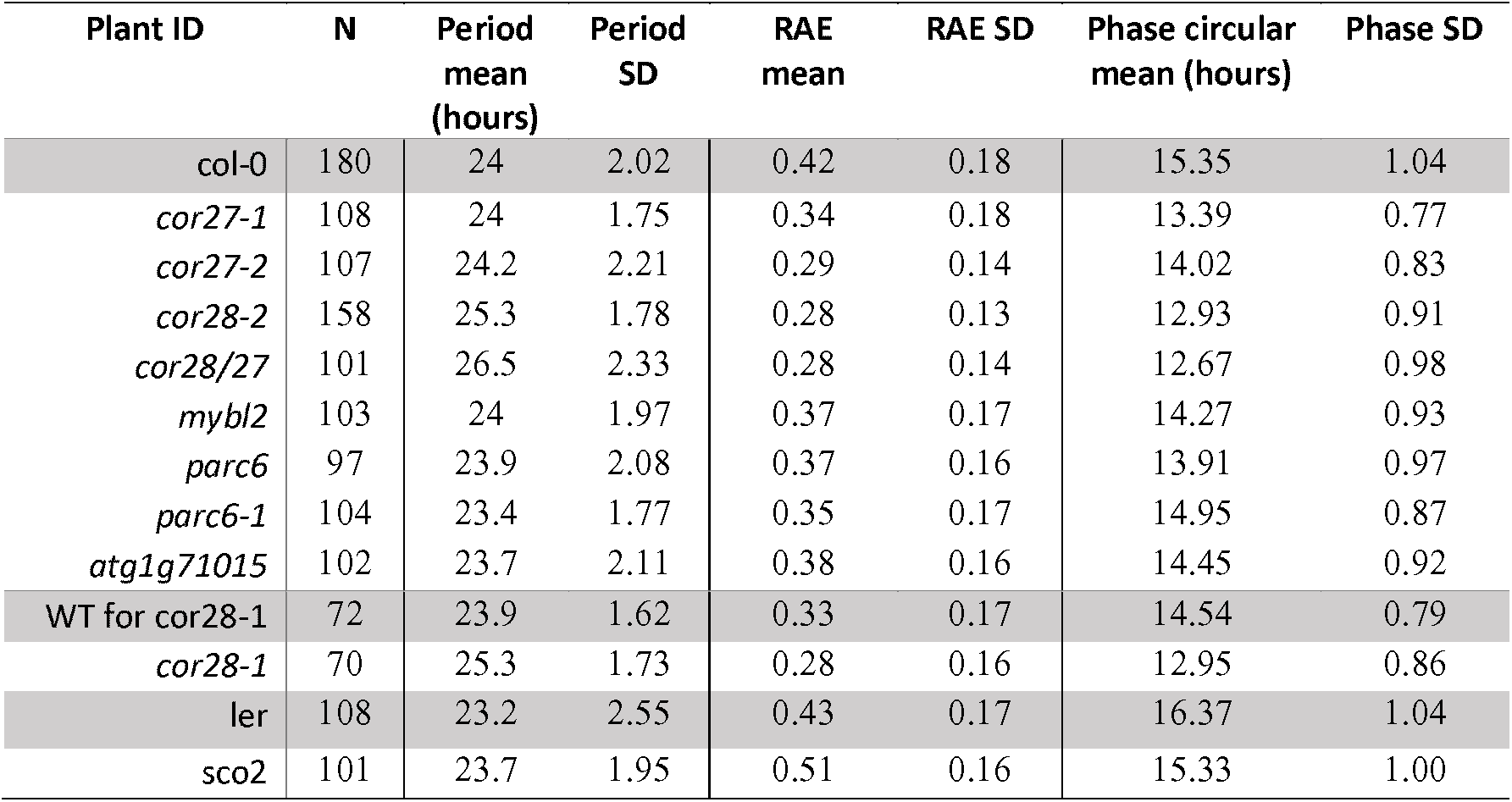
Results from DF screening of GWA candidate mutant lines.

Mutants are listed under their respective WT controls highlighted in grey. Circular means for phase were calculated using the ‘circular’ package in R.

### The phenotypic tails of circadian traits remain largely diverged under lower temperatures

The variation characterized in the first part of this paper was observed at 22°C which was chosen to make the data comparable to several previously published studies of interest. We wanted to test whether the circadian phenotypic diversity observed at 22°C persisted at lower temperatures, closer to the natural Swedish environment. To simplify the dataset, we selected 10 accessions to represent each of the six phenotypic tails as highlighted in Figure 1. We wished to investigate whether reduced temperatures would affect the phenotypic tails equally, drive them further apart or lead to a convergence of their phenotypes. Our null hypothesis was that the phenotypic diversity seen at 22LJ would exist consistently at lower temperatures with no differential effect on the phenotypic tail groups.

The results show that decreasing temperature had a massive overall effect on all three circadian outputs, particularly RAE and phase (accession means can be viewed in Supplementary File 3). Overall, the divergence between the tail groups was largely maintained, although the gap reduced at 10°C (Figure 5a-c). For each trait, a general linear model was fitted to the data in order to test the significance of explanatory factors (Supplementary Tables 15-17).

**Figure 5.**
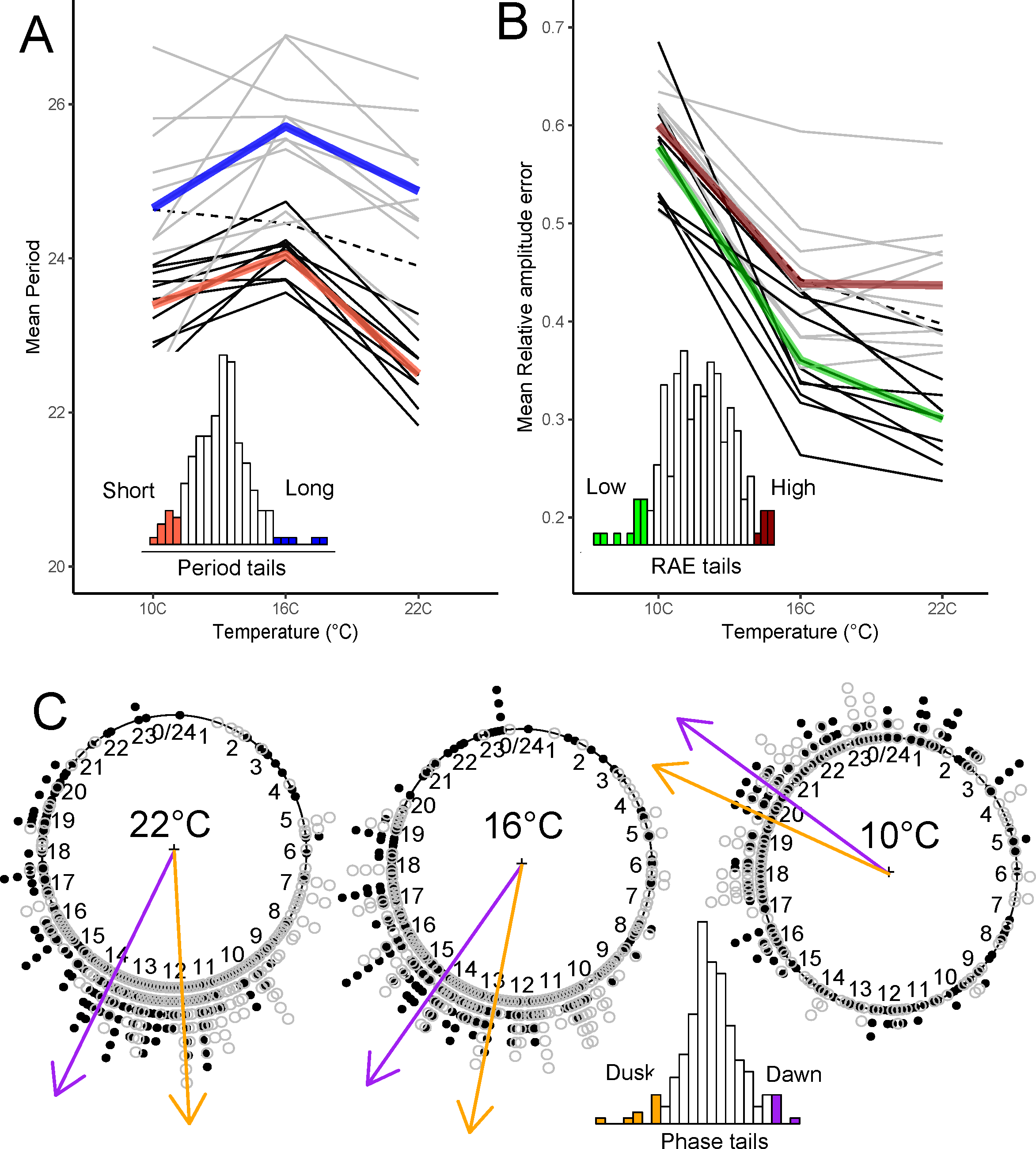
Effects of temperature on circadian rhythms in phenotypic tails. We selected 10 accessions to represent the phenotypic tails of each trait as highlighted in Figure 1. Period, phase and RAE in these accessions were then re-quantified at 10°C, 16°C and 22°C to see if the tails remained diverged at lower temperatures. In Figure A, light grey lines show the mean periods of accessions in the long period tail group with the thick blue line showing the mean period for the whole group. Black lines show the mean periods for accessions in the short period group with the thick red line reflecting the mean for the whole short period group. In Figure B, light grey lines show the mean periods of accessions with high RAE (low robustness) with the thick maroon line indicating the overall high RAE tail mean. The black lines show the mean periods for accessions with low RAE (high robustness). The green line shows the overall low RAE tail mean. In both Figure A and B the dotted line is the mean for Col-0 at each temperature. Figure C shows individual phase estimates from accessions belonging to the dusk-phased (dark filled point) or dawn-phased (open point) tail groups plotted as clock plots relative to 24/0 representing dawn. Tail group phase means are indicated by coloured arrows (dusk tail in orange and dawn tail in purple). Sample size contributing to accession mean at each temperature ranged from 14 to 24.)

For period, membership to the short or long tail groups was the largest explanatory variable and the group means remained clearly distinct across all temperatures. Temperature also had a large overall effect on period, with rhythms at 16°C running much slower than at 22°C. At 10°C periods were again shorter, accompanied by higher RAE (reduced rhythm robustness) (Supplementary Figure 7). The difference in the period temperature responses of the two groups can be seen by the gradients of the thick colored lines in Figure 5A. There was also significant variation between the temperature response of individual accessions within each tail group, especially in the long period group (see thin grey lines in Figure 5A). Interestingly, Col-0 reacted very differently to the Swedish accessions tested, showing an almost linear decrease in period with increasing temperature (dashed line in 5A). For RAE, which we equate to rhythm robustness, we found that temperature had an even greater effect than for period, with rhythms at 10°C showing a marked decrease in robustness (Figure 5B). The tails converge as the temperature decreases, with the low RAE group becoming less rhythmic at a faster rate than the high RAE group.

For phase, decreasing temperature to 10°C caused a large shift of approximately 7.4 hours towards dawn accompanied by increased variability for each accession (Figure 5C). The means of the two phase tail groups remained distinct across the three temperatures and there was no significant difference in their relative change of phase with temperature. However, there was significant variation in responses of individual accessions to temperature within each tail group.

Across all traits we observed an increase in the within-accession variability at lower temperatures indicated by larger standard deviations in period, higher RAE scores and a greater number of rhythms being rejected from Biodare2 analysis as arrhythmic.

## Discussion

Natural variation of any physiological trait is a product of both environmental pressure for local adaption and the available genetic diversity within a population(51). Natural selection acts over evolutionary time on heritable traits which provide a differential fitness cost or advantage. Population migration provides a source of heterozygosity often accompanied by an increase in competitive hybrids, however it also has potential to dilute alleles best adapted to their environmental niche (52). This work provides an example of the interplay between these ecological and evolutionary dynamics.

We measured DF rhythms in 191 naturally occurring *Arabidopsis* accessions and show that circadian phenotypes display considerable variation across the axes of Sweden. This variation does not conform to previously described latitudinal clines in *Arabidopsis*, as we find the longest periods in the South of the country and the shortest periods in the North. We also show a high level of correlation between circadian period, phase and RAE in these accessions; rhythms with longer periods tended to be the most rhythmic with a peak phase just after dusk. The co-variation of these traits with geographical regions reinforce the idea that circadian rhythms are under divergent selection in natural environments. A possible explanation for variation in these traits lies in the massive difference in amplitude of annual temperature and day-length cycles across the length of Sweden. In Northern latitudes an advantage for a dynamic clock with high plasticity could potentially outweigh the advantage of a highly synchronized one with high rhythmicity.

In addition to the geographic variation, periods were also found to be significantly different between two demographic groups in the South. These genetically distinct populations existing within the same geographic area demonstrate the interaction between genotype diversity and adaptation in deciding circadian phenotypes.

We show that circadian diversity assayed by DF is genetically heritable and is associated with several highly significant polymorphisms, two of which (*ELF3* and *COR28*) had previously acknowledged circadian functions. *COR28* is partially redundant with its partner *COR27* and acts both upstream and downstream of the circadian clock(49,50). TDNA insertions in these genes have been shown to lengthen periods, extend flowering time and increase freezing tolerance(49,50). Natural allelic diversity in *COR28* has not previously been described. Here, we show that a set of 16 naturally occurring accessions from southern Sweden have a W58S amino acid substitution which results in a long period comparable to that seen in *cor28* TDNA insert mutants. *COR28* and *COR27* are expressed in a blue-light and temperature dependent manor, which suggests why this gene could be under selection in the Swedish environment. The mechanism through which W58S affects the function of *COR28* remains unclear. COR28 is a small peptide of ∼26kDa and so may not require active transport for nuclear localization(53). No DNA binding domains have been identified in the COR28 sequence, however it has been suggested to regulate *TOC1* and *PRR5* transcription through the formation of protein complexes(49)(50). It is possible that this modification affects the ability of COR28 to form these protein interactions.

We also investigated the effect of temperature on natural circadian variation between accessions with divergent circadian phenotypes. Temperature had a large effect on period and an even greater one on phase and RAE means in both tail groups, indicating a low level of temperature compensation for these outputs. This also shows that the forces governing compensation for period do not act equally on maintaining constant RAE or peak phase. Interestingly, rhythms appeared to be most robust at 22°C, which might not be expected given that the warmest months in Sweden average around 15-17°C. A similar loss of rhythm robustness at lower temperatures has been observed in wheat(23).

Period had a non-linear relationship with temperature in these accessions, lengthening from 22°C to 16°C, before shortening again at 10C. This arrow shaped profile likely reflects two interacting forces at work; 1) between 22°C and 27°C the acceleration of rhythms due to increased rate kinetics and 2) between 22°C and 10°C the balancing forces of circadian temperature compensation. Gould et al. found a similar effect of temperature on period using leaf-movement rhythms at 22°C, 17°C and 12°C, and showed that the cold temperature compensation response works through an independent mechanism to the hot temperature compensation response(20). The profile suggests that temperature compensation is biased towards correction at colder temperatures in these accessions. It is possible that adaptation to a cold climate has selected for a cold compensation response to overcome excessive deceleration of the clock, although we are unable to explain why the rhythms should be even shorter at 10°C than at 16°C or why the shortening of periods are accompanied by a loss of overall rhythmicity. Although divergence between the phenotypic tails decreased at lower temperatures, the groups remained largely separate, reconfirming that these tail phenotypes are due to heritable genotypic differences. This work demonstrates the utility of using DF imaging to analyse natural variation across genetically diverse populations.

## Methods

### Plant material, plate set-up and growth conditions

Seed for the 192 *Arabidopsis* accessions were gas sterilised using 3ml hydrochloric acid in 100ml sodium hypochlorite. We then stratified seed in 1ml of sterile water for 2 days at 4°C before plating. Clear 96 well plates with flat bottomed wells (Thermo Scientific, cat no: 10287631) were filled with 250ul of Murashige and Skoog agar media (1.5% Agar, pH5.8, no sucrose). Approximately 20 seeds were added to the plates following a randomized Alpha-lattice design (see Extended Methods) and left to dry until all residual water had evaporated. Clear microplate lids (Thermo Scientific, cat no: 10334311) were secured to each plate with micropore tape ensuring that the lid was raised slightly from the top of the wells to allow condensation to circulate freely. Plates were returned to the fridge for 2 more days before transfer to the growth cabinet set at 12:12 L:D cycle at 22°C under approximately 200 µmol m^-2^ s^-1^ white light for 14 days. Prior to the temperature response experiments, accessions from the 191 phenotyping screen were bulked for seed (see Extended Methods). Plates were all grown under 12:12 L:D at 22°C light for 10 days and were then transferred to the temperatures in which they would be imaged for 4 days.

Note: For the temperature and mutant screening experiments, the corner wells of the 96-well plates were not used in the designs as we found that corners had significant effects on circadian rhythms in the 191 dataset; potentially due to these wells drying out more quickly (Supplementary Figure 8).

### Imaging conditions

Delayed fluorescence imaging was carried out using Lumo Reteiga CCD cameras (QImaging, Canada) housed in two temperature-controlled cabinets. Six 96-well plates can be imaged simultaneously in each cabinet. Imaging conditions were exactly those described for constant light (L:L) imaging of *Brassica* leaves in Rees et al 2019(23). Images were captured every hour with a 1-minute exposure time.

### FIJI ROI selection and circadian analysis

Image stacks were imported into FIJI (54)and regions of interest (ROI) were selected using a semi-automated approach (full details in Extended methods; macro and user guide in Supplementary files 4 and 5). Measurements for integrated density were taken for these regions across the stack using the Multi-measure plugin. The time-series was converted to decimalized time relative to hours elapsed from entrained dawn (T0) and was cropped to 150h to standardize between experiments. Period estimation was done using the online platform Biodare2 applying the FFT-NLLS algorithm on a data window of 24-120h with expected periods set to 18-34h. Raw data was baseline and amplitude de-trended prior to analysis. Manual inspection of resulting periods ensured that all arrhythmic traces were excluded from further analysis.

### Data analysis and REML

We filtered period estimates from Biodare2 to exclude rhythms with RAE>0.65 and period >28h to improve the precision of mean period predictions. We next used restricted maximum likelihood (REML) to fit a linear mixed model to the 191 accession dataset and thus obtain accession means for period and RAE which were adjusted for the effects of cabinet and experimental run (see details in Extended Methods). An additional step was required to calculate phase means for each accession as the raw phase data is circular and relative to dawn (0) with a full circle representing 24 hours. Phase data was analysed with a circular regression model using the Genstat RCIRCULAR procedure to obtain accession mean phases adjusted for cabinet and run effects(55).

For Temperature experiments, no period or RAE cut-offs were imposed as we predicted an increase in both these variables with lower temperatures. We fitted linear models to the period and RAE data and identified the contribution of each component by analysis of variance (Supplementary Tables 12-14). The above data analysis was done using Genstat 18^th^ edition.

Map figures (Figure 2 and 4) were created using the ggmaps package in R using Google maps (2018)(56).

### Principal component analysis

The PCA for the 191 Swedish accessions in Figure 2E was performed using a compressed mixed linear model run in R implementing the GAPIT package(57,58). Genotype data was downloaded from the 1001 genomes dataset filtered for bialleleic SNPs with >5% MAF (minor allele frequency). The first two PCs were used to split the accessions into the categories in Figure 2D and F.

### GWAS

Genome wide association mapping was conducted using an Accelerated Mixed Model (AMM) via the online web application GWA-Portal(45). Accession means estimated from REML were used for period, phase and RAE with no further transformation necessary. Genotype data used was from the full-sequence 1001 genomes dataset. Results were filtered for the top 10,000-log10(P) values and then for hits with >5% MAF. Comparison to results from other models confirmed the major peaks and can be viewed in Supplementary Figure 6.

### Mutant screening

T-DNA and EMS mutants were obtained from the Nottingham *Arabidopsis* Stock centre and primers designed using the iSect tool from the SIGnAL website. In addition, seed for *cor28-2, cor27-1* and *cor28-2/cor27-1* was kindly donated from Hongtao Liu’s laboratory (Shanghai, China) where these mutants have previously been identified as long-period using leaf-movement and ProCCR2:LUC constructs(49). *Cor28-2* is a null mutant and *cor28-1, cor27-1* and *cor27-2* are all knockdown mutants. Gene candidates, mutant IDs and primers used can be viewed in Supplementary File 2. All lines were genotyped to confirm homozygosity by PCR and gel electrophoresis prior to seed bulking other than *sco2* which was genotyped by sanger sequencing to reveal the premature stop. We were unable to obtain or confirm homozygous mutants for *dnf* and *fab1c* and therefore these lines were not DF screened (Supplementary File 2). SALK lines for *elf3* were not obtained as the *elf3-1* mutant has been previously confirmed in Box et al. 2015 as having a phenotype of reduced robustness using delayed fluorescence imaging(59). WT control lines; Col-0 and Ler were grown for seed in parallel to standardize seed quality. For *cor28-1*, a homozygous WT control was derived by segregation of the original heterozygous TDNA line was used as a control. Five imaging experiments using a single camera and cabinet were carried out with each line present in at least three experiments. No mixed model was applied to the mutant screening because only one camera was used.

## Supporting information

Supplementary File 1. phenotype files from 191 dataset

Supplementary File 2. Gene candidates for DF phenotyping

Supplementary File 5. Selecting ROI for DF imaging using multiple 96 well plates

Supplementary File 3. Line means and SD for temperature experiments

Supplementary File 4. generate_all_rois_96_plate

## Acknowledgements

We thank Paul W. Goedhart (Wageningen University) for advice about circular phase statistics using the RCIRCULAR procedure and Grant Calder (John Innes Centre) for his assistance developing the 96-well ROI selection using FIJI. We thank Susan Duncan for help and advice with the flowering time investigation.

We also thank the Dean Research group (JIC) for providing the 191 Swedish accessions and the Liu Research group (Shanghai Institutes for Biological Sciences) for providing the *cor28-2, cor27-1* and *cor27-1/28-2* mutants. We also thank Ewan Holmes and Georgia Scutter for help with seed harvesting.

## References

1. Harmer SL, Hogenesch JB, Straume M, Chang HS, Han B, Zhu T, et al. Orchestrated transcription of key pathways in Arabidopsis by the circadian clock. Science. 2000 Dec 15;290(5499):2110–3.

2. Covington MF, Maloof JN, Straume M, Kay SA, Harmer SL. Global transcriptome analysis reveals circadian regulation of key pathways in plant growth and development. Genome Biol. 2008;9(8):R130.

3. Dodd AN, Salathia N, Hall A, Kévei E, Tóth R, Nagy F, et al. Plant circadian clocks increase photosynthesis, growth, survival, and competitive advantage. Science. 2005;309(5734):630–3.

4. Green RM, Tingay S, Wang Z-Y, Tobin EM. Circadian rhythms confer a higher level of fitness to Arabidopsis plants. Plant Physiol. 2002 Jun 1;129(2):576–84.

5. Ingle RA, Stoker C, Stone W, Adams N, Smith R, Grant M, et al. Jasmonate signalling drives time-of-day differences in susceptibility of Arabidopsis to the fungal pathogen Botrytis cinerea. Plant J. 2015 Dec;84(5):937–48.

6. Michael TP, Salomé PA, Yu HJ, Spencer TR, Sharp EL, McPeek MA, et al. Enhanced Fitness Conferred by Naturally Occurring Variation in the Circadian Clock. Science (80-). 2003;302(5647):1049–53.

7. Hernando CE, Romanowski A, Yanovsky MJ. Transcriptional and post-transcriptional control of the plant circadian gene regulatory network. Biochim Biophys Acta - Gene Regul Mech. 2017 Jan 1;1860(1):84–94.

8. Alabadi D, Oyama T, Yanovsky MJ, Harmon FG, Más P, Kay SA. Reciprocal Regulation Between TOC1 and LHY/CCA1 Within the Arabidopsis Circadian Clock. Science (80-). 2001 Aug 3;293(5531):880–3.

9. Kinmonth-Schultz HA, Golembeski GS, Imaizumi T. Circadian clock-regulated physiological outputs: dynamic responses in nature. Semin Cell Dev Biol. 2013 May;24(5):407–13.

10. Suárez-López P, Wheatley K, Robson F, Onouchi H, Valverde F, Coupland G. CONSTANS mediates between the circadian clock and the control of flowering in Arabidopsis. Nature. 2001 Apr 26;410(6832):1116–20.

11. Johanson U, West J, Lister C, Michaels S, Amasino R, Dean C. Molecular analysis of FRIGIDA, a major determinant of natural variation in Arabidopsis flowering time. Science. 2000 Oct 13;290(5490):344–7.

12. Bastow R, Mylne JS, Lister C, Lippman Z, Martienssen RA, Dean C. Vernalization requires epigenetic silencing of FLC by histone methylation. Nature. 2004 Jan 8;427(6970):164–7.

13. James AB, Syed NH, Bordage S, Marshall J, Nimmo GA, Jenkins GI, et al. Alternative splicing mediates responses of the Arabidopsis circadian clock to temperature changes. Plant Cell. 2012 Mar;24(3):961–81.

14. Pittendrigh CS, Minis DH. The Entrainment of Circadian Oscillations by Light and Their Role as Photoperiodic Clocks. Am Nat. 1964 Sep 15;98(902):261–94.

15. Aschoff J. Circadian rhythms: influences of internal and external factors on the period measured in constant conditions. Z Tierpsychol. 1979 Mar;49(3):225–49.

16. Yanovsky MJ, Kay SA. Molecular basis of seasonal time measurement in Arabidopsis. Nature. 2002 Sep 19;419(6904):308–12.

17. Más P, Alabadí D, Yanovsky MJ, Oyama T, Kay SA. Dual role of TOC1 in the control of circadian and photomorphogenic responses in Arabidopsis. Plant Cell. 2003 Jan 1;15(1):223–36.

18. Edwards KD, Lynn JR, Gyula P, Nagy F, Millar AJ. Natural allelic variation in the temperature-compensation mechanisms of the Arabidopsis thaliana circadian clock. Genetics. 2005 May;170(1):387–400.

19. Salome PA, McClung CR. PSEUDO-RESPONSE REGULATOR 7 and 9 Are Partially Redundant Genes Essential for the Temperature Responsiveness of the Arabidopsis Circadian Clock. PLANT CELL ONLINE. 2005 Feb 18;17(3):791–803.

20. Gould PD, Locke JCW, Larue C, Southern MM, Davis SJ, Hanano S, et al. The Molecular Basis of Temperature Compensation in the Arabidopsis Circadian Clock. PLANT CELL ONLINE. 2006 May 1;18(5):1177–87.

21. Colin Pittendrigh BS, by N Harvey CE. On temperature independence in the clock-system controlling emergence Time in Drosophila. Vol. 102, Proc. Soc. Exptl. Biol. Med. 1954.

22. Kusakina J, Gould PD, Hall A. A fast circadian clock at high temperatures is a conserved feature across Arabidopsis accessions and likely to be important for vegetative yield. Plant Cell Environ. 2014 Feb;37(2):327–40.

23. Rees H, Duncan S, Gould P, Wells R, Greenwood M, Brabbs T, et al. A high-throughput delayed fluorescence method reveals underlying differences in the control of circadian rhythms in Triticum aestivum and Brassica napus. Plant Methods. 2019 May;15(1):51.

24. Allemand R, David JR. The Circadian Rhythm of Oviposition in Drosophila melanogaster: A Genetic Latitudinal Cline in Wild Populations. Vol. 33, Jap. J. Pharmac. Princeton Univ. Press; 1974.

25. Lankinen P. Comparative a Geographical variation in circadian eclosion rhythm and photoperiodic adult diapause in Drosophila littorMis. Vol. 159, Journal of Sensory. 1986.

26. Swarup K, Alonso-Blanco C, Lynn JR, Michaels SD, Amasino RM, Koornneef M, et al. Natural allelic variation identifies new genes in the Arabidopsis circadian system. Plant J. 1999;20(1):67–77.

27. de Montaigu A, Giakountis A, Rubin M, Tóth R, Cremer F, Sokolova V, et al. Natural diversity in daily rhythms of gene expression contributes to phenotypic variation. Proc Natl Acad Sci U S A. 2015 Jan 20;112(3):905–10.

28. Anwer MU, Boikoglou E, Herrero E, Hallstein M, Davis AM, Velikkakam James G, et al. Natural variation reveals that intracellular distribution of ELF3 protein is associated with function in the circadian clock. Elife. 2014 May 27;3.

29. Müller NA, Wijnen CL, Srinivasan A, Ryngajllo M, Ofner I, Lin T, et al. Domestication selected for deceleration of the circadian clock in cultivated tomato. Nat Genet. 2016 Jan 16;48(1):89–93.

30. Greenham K, Lou P, Puzey JR, Kumar G, Arnevik C, Farid H, et al. Geographic Variation of Plant Circadian Clock Function in Natural and Agricultural Settings. J Biol Rhythms. 2017 Feb 26;32(1):26–34.

31. Shindo C, Aranzana MJ, Lister C, Baxter C, Nicholls C, Nordborg M, et al. Role of FRIGIDA and FLOWERING LOCUS C in determining variation in flowering time of Arabidopsis. Plant Physiol. 2005 Jun;138(2):1163–73.

32. Bloomer RH, Dean C. Fine-tuning timing: natural variation informs the mechanistic basis of the switch to flowering in Arabidopsis thaliana. J Exp Bot. 2017 Nov 28;68(20):5439–52.

33. Nordborg M, Hu TT, Ishino Y, Jhaveri J, Toomajian C, Zheng H, et al. The Pattern of Polymorphism in Arabidopsis thaliana. Mitchell-Olds T, editor. PLoS Biol. 2005 May 24;3(7):e196.

34. Horton MW, Hancock AM, Huang YS, Toomajian C, Atwell S, Auton A, et al. Genome-wide patterns of genetic variation in worldwide Arabidopsis thaliana accessions from the RegMap panel. Nat Genet. 2012 Jan 8;44(2):212–6.

35. Long Q, Rabanal FA, Meng D, Huber CD, Farlow A, Platzer A, et al. Massive genomic variation and strong selection in Arabidopsis thaliana lines from Sweden. Nat Genet. 2013 Aug 23;45(8):884–90.

36. Huber CD, Nordborg M, Hermisson J, Hellmann I. Keeping it local: evidence for positive selection in Swedish Arabidopsis thaliana. Mol Biol Evol. 2014 Nov;31(11):3026–39.

37. Alonso-Blanco C, Andrade J, Becker C, Bemm F, Bergelson J, Borgwardt KM, et al. 1,135 Genomes Reveal the Global Pattern of Polymorphism in Arabidopsis thaliana. Cell. 2016 Jul 14;166(2):481–91.

38. Exposito-Alonso M, Vasseur F, Ding W, Wang G, Burbano HA, Weigel D. Genomic basis and evolutionary potential for extreme drought adaptation in Arabidopsis thaliana Europe PMC Funders Group. Nat Ecol Evol. 2018;2(2):352–8.

39. Weigel D, Mott R. The 1001 Genomes Project for Arabidopsis thaliana. Genome Biol. 2009;10(5):107.

40. Kerdaffrec E, Filiault DL, Korte A, Sasaki E, Nizhynska V, Seren Ü, et al. Multiple ?alleles at a single locus control seed dormancy in Swedish Arabidopsis. Elife. 2016;5.

41. Sasaki E, Zhang P, Atwell S, Meng D, Nordborg M. “Missing” G x E Variation Controls Flowering Time in Arabidopsis thaliana. Gibson G, editor. PLOS Genet. 2015 Oct 16;11(10):e1005597.

42. Horton MW, Willems G, Sasaki E, Koornneef M, Nordborg M. The genetic architecture of freezing tolerance varies across the range of *Arabidopsis thaliana*. Plant Cell Environ. 2016 Nov 1;39(11):2570–9.

43. Gould PD, Diaz P, Hogben C, Kusakina J, Salem R, Hartwell J, et al. Delayed fluorescence as a universal tool for the measurement of circadian rhythms in higher plants. Plant J. 2009 Jun;58(5):893–901.

44. Li Y, Huang Y, Bergelson J, Nordborg M, Borevitz JO. Association mapping of local climate-sensitive quantitative trait loci in Arabidopsis thaliana PLANT BIOLOGY. 2010;7(49):21199–204.

45. Seren Ü. GWA-Portal: Genome-Wide Association Studies Made Easy. In Humana Press, New York, NY; 2018. p. 303–19.

46. Proietti S, Caarls L, Coolen S, Van Pelt JA, Van Wees SCM, Pieterse CMJ. Genome-wide association study reveals novel players in defense hormone crosstalk in Arabidopsis. Plant Cell Environ. 2018;41(10):2342–56.

47. van Rooijen R, Aarts MGM, Harbinson J. Natural Genetic Variation for Acclimation of Photosynthetic Light Use Efficiency to Growth Irradiance in Arabidopsis. Plant Physiol. 2015;167(4):1412–29.

48. Kooke R, Kruijer W, Bours R, Becker F, Kuhn A, van de Geest H, et al. Genome-Wide Association Mapping and Genomic Prediction Elucidate the Genetic Architecture of Morphological Traits in Arabidopsis. Plant Physiol. 2016;170(4):2187–203.

49. Li X, Ma D, Lu SX, Hu X, Huang R, Liang T, et al. Blue Light- and Low Temperature-Regulated COR27 and COR28 Play Roles in the Arabidopsis Circadian Clock. Plant Cell. 2016;28(11):2755–69.

50. Wang P, Cui X, Zhao C, Shi L, Zhang G, Sun F, et al. COR27 and COR28 encode nighttime repressors integrating Arabidopsis circadian clock and cold response. Vol. 59, Journal of Integrative Plant Biology. 2017. p. 78–85.

51. Andrews CA. Natural Selection, Genetic Drift, and Gene Flow Do Not Act in Isolation in Natural Populations. Nat Educ Knowl. 2010;1(10):5.

52. Bourne EC, Bocedi G, Travis JMJ, Pakeman RJ, Brooker RW, Schiffers K. Between migration load and evolutionary rescue: Dispersal, adaptation and the response of spatially structured populations to environmental change. Proc R Soc B Biol Sci. 2014;281(1778).

53. Hicks GR. Nuclear Import of Plant Proteins. In: Madame Curie Bioscience Database [Internet]. Austin (TX): Landes Bioscience; 2013.

54. Schindelin J, Arganda-Carreras I, Frise E, Kaynig V, Longair M, Pietzsch T, et al. Fiji: an open-source platform for biological-image analysis. Nat Methods. 2012 Jul 1;9(7):676–82.

55. Fisher NI, Lee AJ. Regression Models for an Angular Response. Biometrics. 1992 Sep;48(3):665.

56. Kahle D, Wickham H. ggmap: Spatial Visualization with ggplot2. R J. 2013;5(1):144–161.

57. Lipka AE, Tian F, Wang Q, Peiffer J, Li M, Bradbury PJ, et al. GAPIT: Genome association and prediction integrated tool. Bioinformatics. 2012;28(18):2397–9.

58. Zhang Z, Ersoz E, Lai CQ, Todhunter RJ, Tiwari HK, Gore MA, et al. Mixed linear model approach adapted for genome-wide association studies. Nat Genet. 2010;42(4):355–60.

59. Box MS, Huang BE, Domijan M, Jaeger KE, Khattak AK, Yoo SJ, et al. ELF3 Controls Thermoresponsive Growth in Arabidopsis. Curr Biol. 2015 Jan 19;25(2):194–9.

60. Thorsen S. timeanddate.com [Internet]. [cited 2019 Jun 7]. Available from: https://www.timeanddate.com/information/copyright.html

