## Extended Methods JB 29-5-19 for "Circadian diversity in Swedish *Arabidopsis* accessions is associated with naturally occurring genetic variation in *COR28*"

Bulking of seed for temperature experiments

Seeds were sown directly onto soil (JIC Arabidopsis mix: Levington F2 compost plus grit) and grown for 2 weeks in long day (16h:8h) conditions at 22°C before a 12-week vernalisation at 4°C. Plants were then moved back to the long day conditions and left to grow before bagging and seed collection. The 45 accessions from the phenotypic tails were prepared for imaging in exactly the same way as for the 191 accession experiments, following a new alpha-lattice design.

Experimental designs

For circadian phenotyping of 191 accessions: The Alpha module in Gendex (<http://www.designcomputing.net/>) was used to create an alpha-lattice design with the following arguments: *v*=192 (number of treatments *v=ks)*, *r*=18 (number of replicates), *k*=24 block size (1/4 of a 96 well plate) and *s*=8 (number of blocks per replicate). Each experiment consisted of 12 96-well plates split between two imaging cabinets and experiments were done in three replicates. Two 96-well plates held a complete set of 191 accessions plus 1 Columbia control which were replicated 18 times in total.

For temperature response experiments: Three experiments were run with each temperature (10°C, 16°C and 22°C) being tested in two replicates, once in each cabinet. We created an alpha-lattice design for 6 plates in 1 cabinet which we used in each experiment. The arguments in this case were: v=48, r=12, k=16 (1/6 of a 96 well plate) and s=3. We manually edited the design so that the corners of each plate were empty (as the design has 6 blanks per plate and we routinely lost corner wells in the phenotyping experiments, see Supplementary Figure 8). This design allowed each of the 44 representative accessions (plus a Columbia control) to be imaged 24 times in each temperature condition.

For mutant screening we ran three experiments of 3+3+6 plates. The alpha-lattice design was for three plates which was then replicated in each run (v=12, r=4, k=4, s=3). We screened 11 lines and a Col-0 control with a total of 96 replicates of each line.

Imaging conditions

Images were captured every hour using Lumo Reteiga CCD cameras fitted with a Xenon 0.95/25mm lens from Schneider-Kreuznach. Cameras were housed in temperature controlled Sanyo MIR-553 growth cabinets fitted with a red/blue LED rig. The LEDs could be instantly turned on and off through an Arduino Uno microcontroller board responding to a uManager software interface. Camera properties were kept the same in each experiment (Binning=4, Gain=1, Readout-Rate=0.650195MHz 16 bit) and camera exposure was initiated 500ms after the lights were turned off. An image was captured after a 1-minute exposure following 59 minutes in the light.

FIJI ROI selection

The add-in Circle-tool macro was set to an appropriate radius for the wells in the image and then the top-left (A1) well was selected from each plate in the image from 1 to 6. These ‘plate identifier’ ROI’s were labelled in the ROI manager. A customised macro (Supplementary Files 4 and 5) was then run which draws ROI in a 96 well format using the A1 wells as a reference. ROIs can be translated or nudged into position until the whole well is centrally enclosed within the selection. Measurements for integrated density were taken for these regions across the stack using the Multi-measure plugin.

Data analysis and REML

Means for period, phase, amplitude and RAE for circadian phenotyping of 191 accessions were obtained as follows. Individual wells with RAE values >0.65 and period estimates >28h were removed from further analysis because the accuracy of predictions was reduced above these cut-offs. Amplitude values were log_10_ transformed to improve normality for linear mixed modeling. The mean period values for each well in all 96-well plates revealed significant effects of each of the 4 corner wells (Supplementary Figure 8) and so these wells were also excluded from further analysis for all traits. A linear mixed model was then fitted to the data for each trait using the REML command in Genstat. The model for each trait took the format:

$$y_{ijklm}=\mu+c_{l}+e_{k}+{c.e}_{lk}+r_{i}+b_{m}+a_{j}+ \varepsilon_{\begin{aligned} ijklm \\ \end{aligned}}$$

where $y_{ijklm}$ is the trait as a dependent variable, $c_{l}$ is the cabinet fixed effect (l = 1,2), $e_{k}$ is the experimental run fixed effect, $a_{j}$ is the accession ID as a random effect, $r_{i}$ is the replicate random effect, $b_{m}$ is the block random effect and $\varepsilon_{ijklm}$ is the residual error. A log10 transformation was used to analyse amplitude (Log10Amplitude) in order to normalise residual errors and make them independent of fitted values.

An additional step was required to calculate average phase means for each accession as the raw phase data is relative to dawn (0) and is circular with a full circle representing 24 hours. This was first converted into degrees with a full circle representing 360°. The Genstat RCIRCULAR procedure was then used to estimate accession means adjusted for cabinet and run effects.

To assess the effect of accession ID on each trait we performed a likelihood ratio test for the model including accession ID against a model in which accession was not included as a random effect. We also calculated the Akaike information criteria (AIC) for the same pairs of models, which fully supported the conclusions of the likelihood ratio tests (Supplementary Tables 4-7). For period, including accession ID in the mixed model had a statistically significant effect on period (χ^2^ (1)=454.36, *p*<0.0001). RAE was also significantly affected by adding accession ID to the model (χ^2^ (1)= 65.396, *p*<0.0001). Accession ID did not have a significant effect on Log10Amplitude. With this in mind, we chose only the tails of period, phase and RAE to take forward for the temperature experiments.

For Temperature experiments:

No period or RAE cut-offs were imposed on data in the experiment with three temperatures as we predicted an increase in both of these variables with lower temperatures. We fitted general linear models (LMs) to the period and RAE data and tested the contribution of each component to the trait by analysis of variance (Supplementary Tables 15-17). The models were as follows:

$$y_{jlno}={\mu+c}_{l}+t_{n}+p_{o}+{c.p}_{lo}+{t.p}_{no}+{p.a}_{oj}+ {c.p.a}_{loj}+{t.p.a}_{noj}+\varepsilon_{jlno}$$

where $y_{jlno}$ is the trait as a dependent variable, $c_{l}$ is the cabinet fixed effect, $t_{n}$ is the temperature fixed effect, $p_{o}$ is the phenotype tail $a_{j}$ is the accession ID within each phenotype tail set and $\varepsilon_{jlno}$ is the residual error.

To calculate effects of temperature on the circular means of the phase tail accessions we performed circular regression using the RCIRCULAR procedure as for the 191 accession dataset. As before, we sequentially removed model terms to test the contributions of each component to the fit of the model. Supplementary File 3 shows mean traits for each accession averaged across the two imaging cabinets.
