## Supplementary Figures 1-8 and Tables 1-17 for "Circadian diversity in Swedish *Arabidopsis* accessions is associated with naturally occurring genetic variation in *COR28*"

Supplementary Figures and Tables:

| | |
|---|---|
| Supplementary Figure 1 | Genstat output, residual Plots to check for normality and outliers in period data from 191 accessions (191 accession data) |
| Supplementary Figure 2 | Genstat output, residual Plots to check for normality and outliers in RAE data from 191 accessions (191 accession data) |
| Supplementary Figure 3 | Genstat output, residual Plots to check for normality and outliers in Log10Amp data from 191 accessions |
| Supplementary Figure 4. | Correlations between REML adjusted accession means for Period, Phase and RAE for each accession in the 191 accession dataset. |
| Supplementary Figure 5 | Period correlation with latitude and longitude (191 accession data) |
| Supplementary Figure 6 | Manhattan plots and Q-Q plots from all GWA models (191 accession data) |
| Supplementary Figure 7 | Period and RAE correlations in Period tail accessions (Temperature data) |
| Supplementary Figure 8 | Position in 96-well plate affects period estimation-justification for removing these wells from analysis (191 accession data). |
| Supplementary Table 1 | Output from Genstat using the REML directive on 191 period data |
| Supplementary Table 2 | Output from Genstat using the REML directive on 191 RAE data |
| Supplementary Table 3 | Output from Genstat using the REML directive on 191 log10Amplitude data |
| Supplementary Table 4 | Likelihood test for period (191 accession data) |
| Supplementary Table 5 | Likelihood test for phase (191 accession data) |
| Supplementary Table 6 | Likelihood test for RAE (191 accession data) |

| | |
|---|---|
| Supplementary Table 7 | Likelihood test for Log10Amp (191 accession data) |
| Supplementary Table 8 | Accessions in Period Tails for temperature experiments (Temperature data) |
| Supplementary Table 9 | Accessions in Phase Tails for temperature experiments (Temperature data) |
| Supplementary Table 10 | Accessions in RAE Tails for temperature experiments (Temperature data) |
| Supplementary Table 11 | Linear regression with previously published datasets (191 accession data) |
| Supplementary Table 12 | Testing differences in Period (Welch Two Sample t-test) (Mutant validation data) |
| Supplementary Table 13 | Supplementary Table S3: Testing differences in RAE (Welch Two Sample t-test) (Mutant validation data) |
| Supplementary Table 14 | Supplementary Table S4: Testing differences in Phase (Watson's Two-Sample Test of Homogeneity) (Mutant validation data) |
| Supplementary Table 15 | Accumulated analysis of varience table for period with temperature data |
| Supplementary Table 16 | Accumulated analysis of varience table for RAE with temperature data |
| Supplementary Table 17 | Circular regression analysis for Phase with temperature data |

Model Checking: Period, RAE, Log10Amplitude

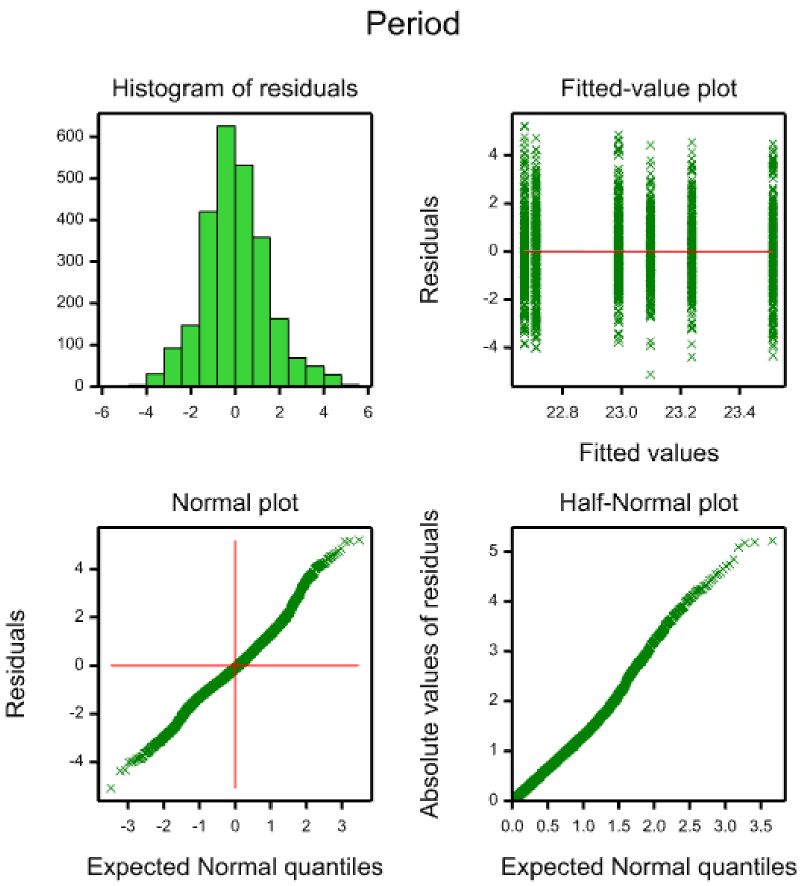


**Supplementary Figure 1.** Plots to examine normality and independence of residuals from the linear mixed model of period in 191 accessions of Arabidopsis thaliana. Output from Genstat 18th edition.

### RAE

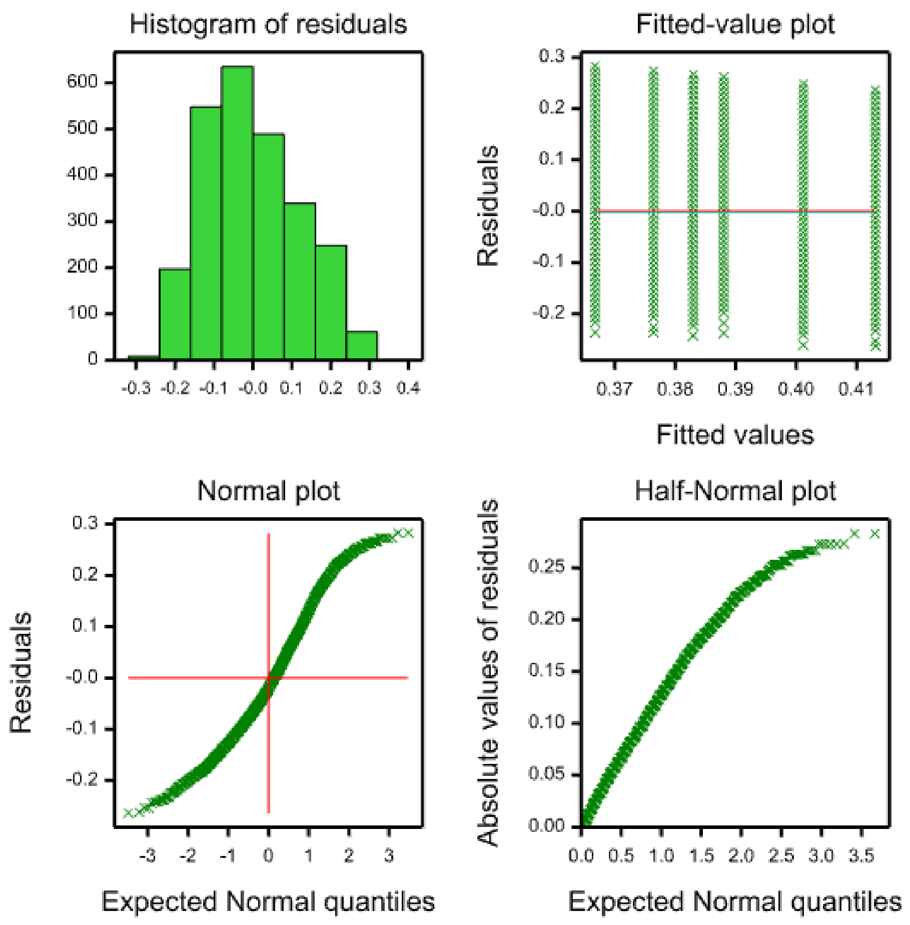


**Supplementary Figure 2.** Plots to examine normality and independence of residuals from the linear mixed model of relative amplitude error in 191 accessions of Arabidopsis thaliana. Output from Genstat 18th edition.

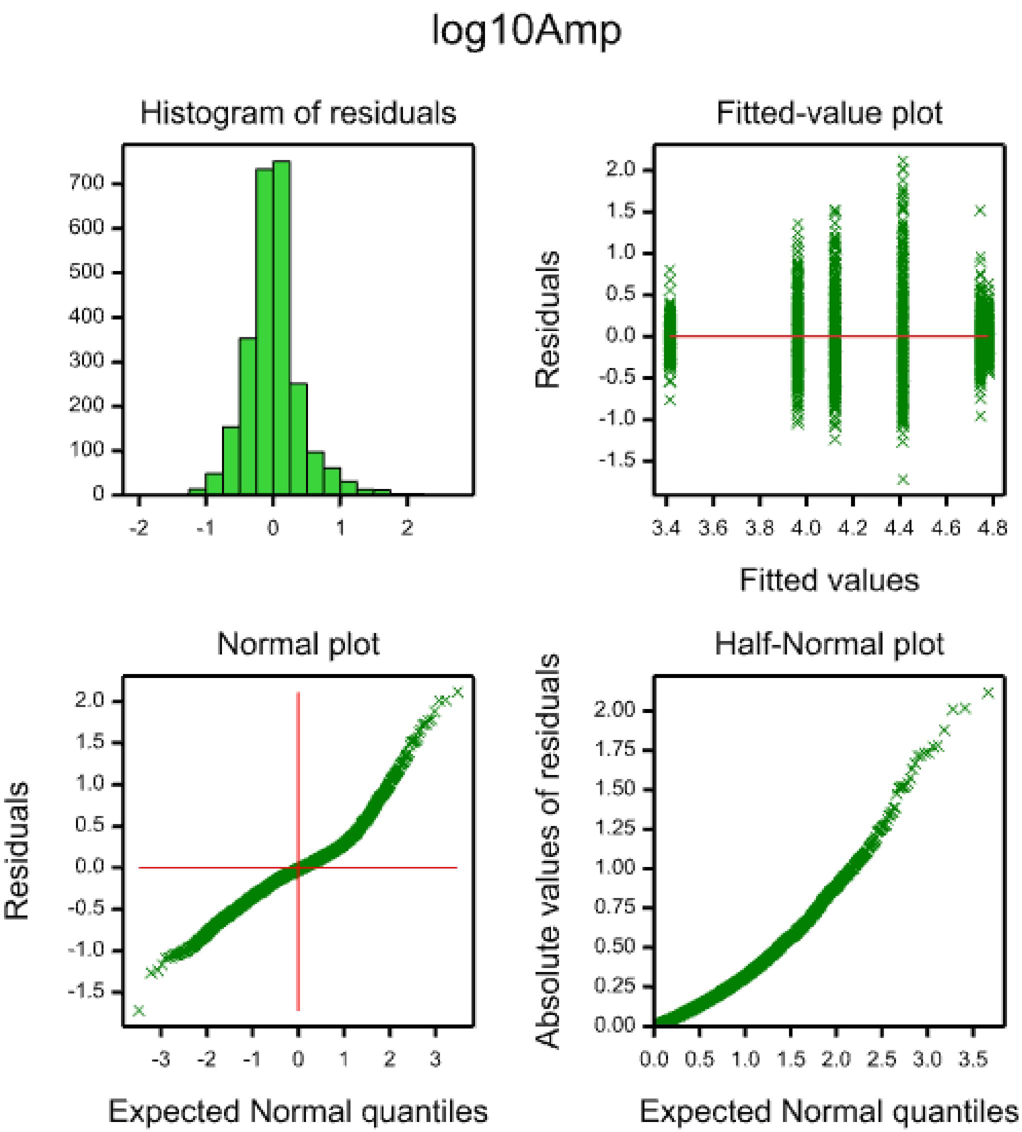


**Supplementary Figure 3.** Plots to examine normality and independence of residuals from the linear mixed model of log10 (amplitude) in 191 accessions of Arabidopsis thaliana. Output from Genstat 18th edition. Note that residuals of a model of untransformed amplitude deviated greatly from the modelling assumptions.

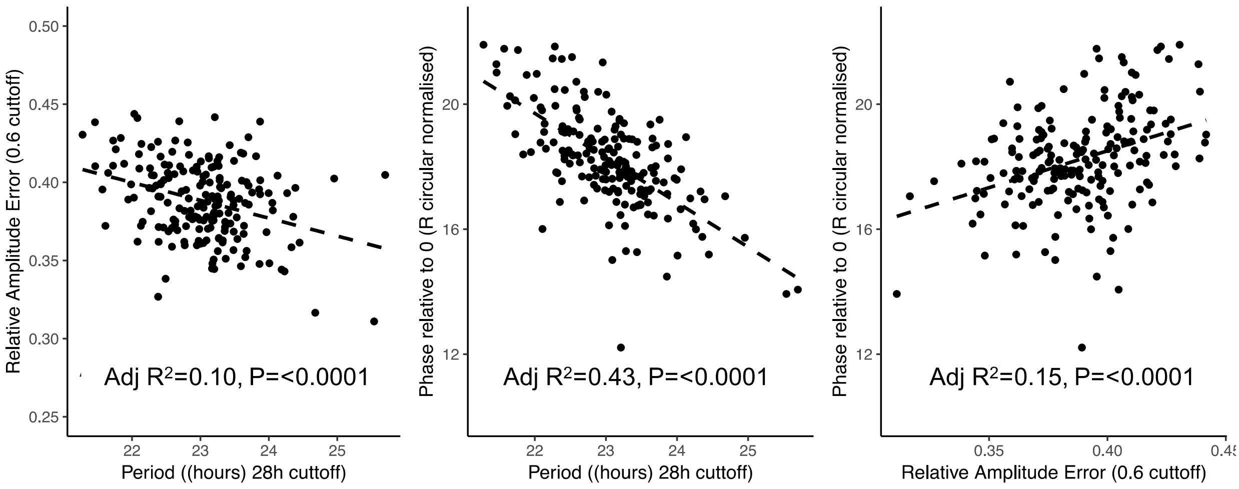


**Supplementary Figure 4.** Correlations between predicted means of period, phase and RAE for 191 accessions of Arabidopsis thaliana, calculated from appropriate linear mixed models using REML. Adjusted R2 and P-values were calculated from linear regression, using an F-test with 1 and 189 degrees of freedom.

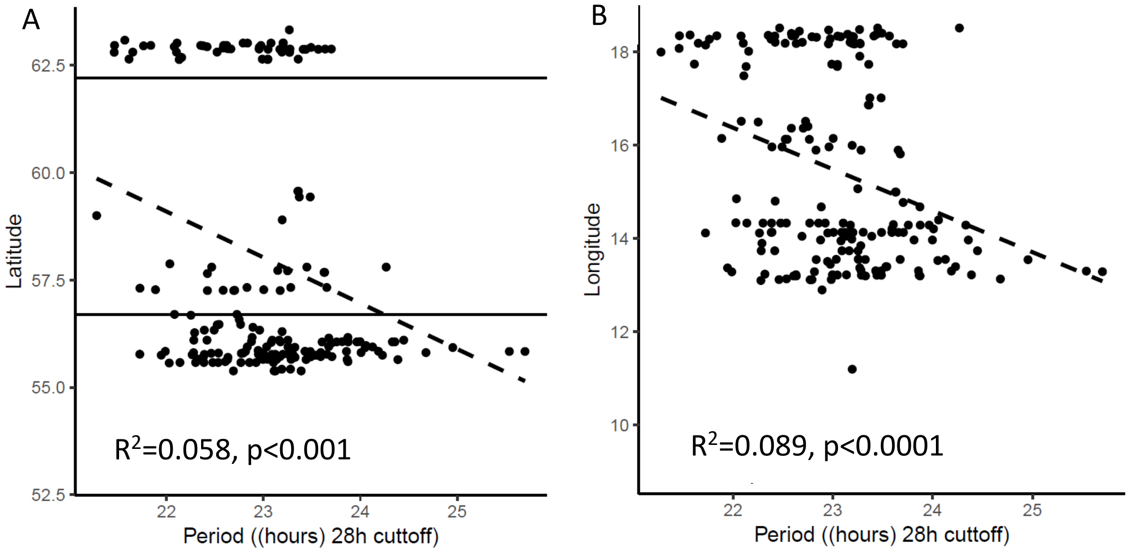


**Supplementary Figure 5.** Correlation of predicted mean period with latitude and longitude for 191 accessions of Arabidopsis thaliana.
Period is significantly correlated with latitude (A). Horizontal lines differentiate between North, Mid and South Sweden. Period is significantly correlated with longitude (B).

**Supplementary Figure 6.** Manhattan plots and respective Q-Q plots from all GWA models (linear, Kruskal-Wallis and AMM) available from GWA-online portal. Results are plotted as negative log-transformed *p* values from associations with the three circadian traits measured in this study: period, RAE and phase. Bonferroni (red dotted line) and Benjamini Hochberg (blue dotted line) are thresholds provided for reference from the GWA output but were not used for significant SNP selection in this study.

### Period

#### Linear model

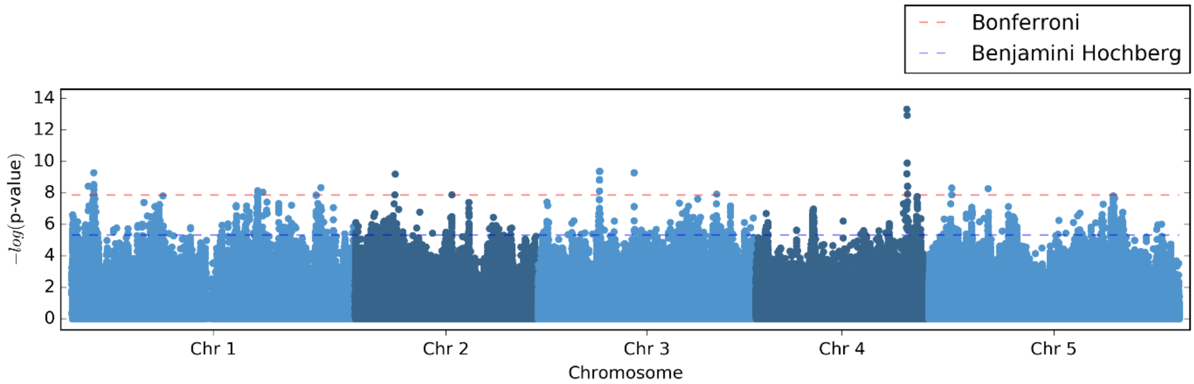


#### Kruskal-Wallis Model

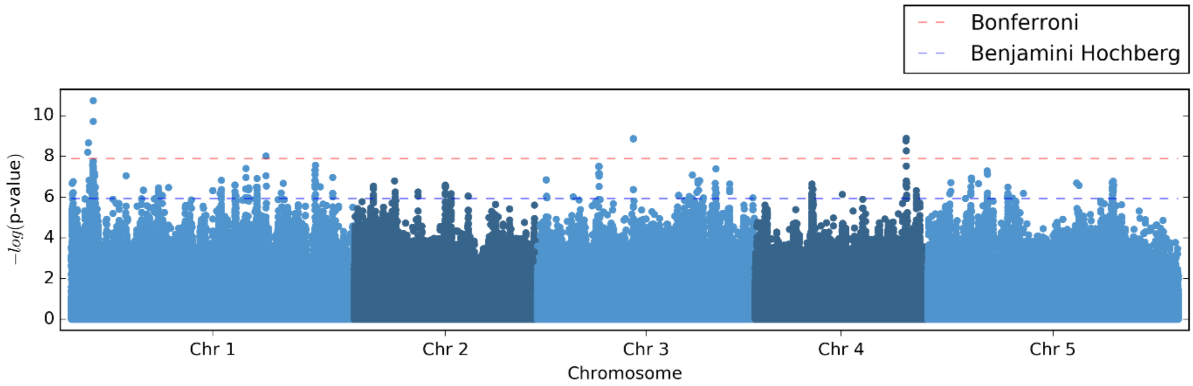


#### Accelerated Mixed Model

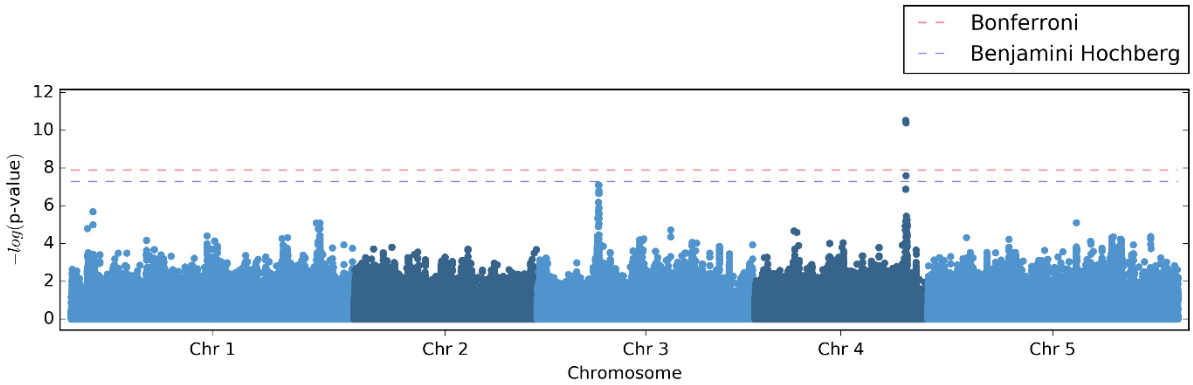

#### Linear model

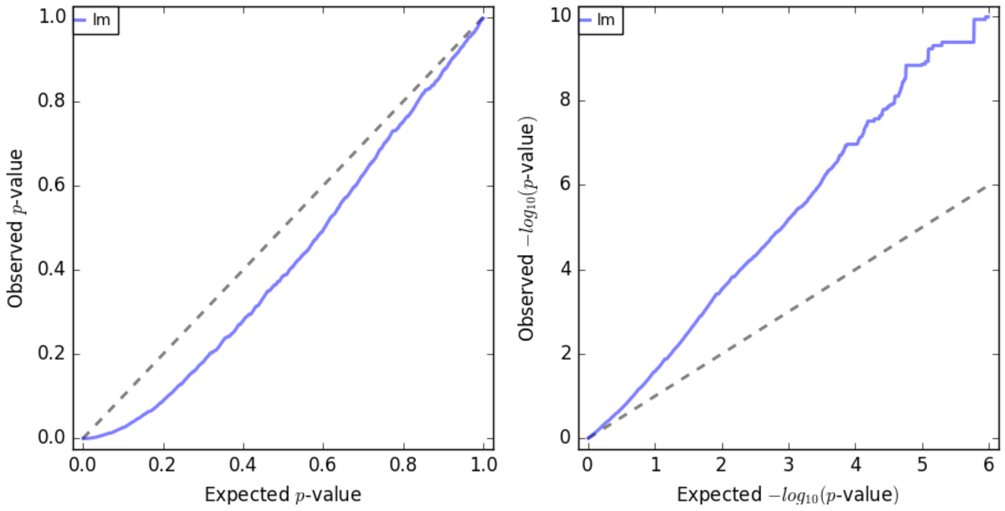


#### Kruskal-Wallis Model

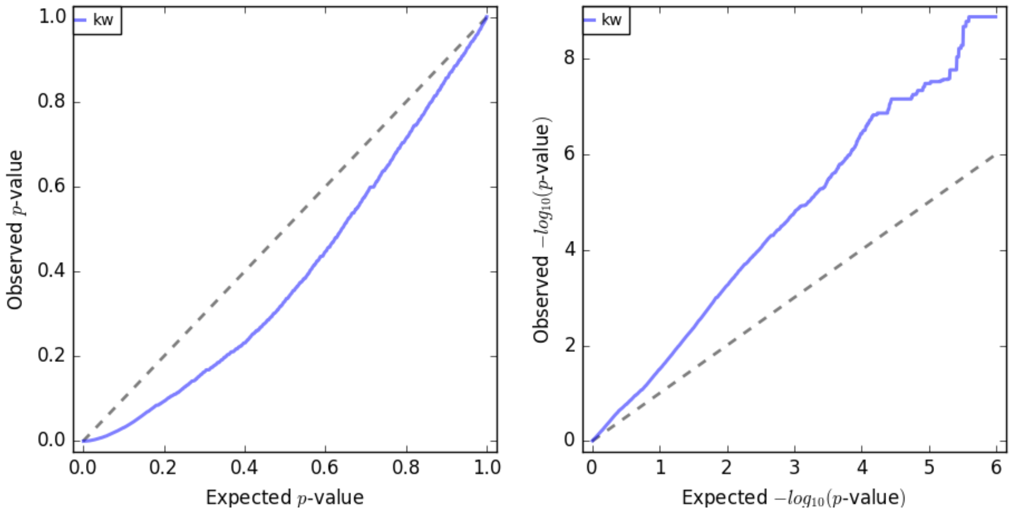


#### Accelerated Mixed Model

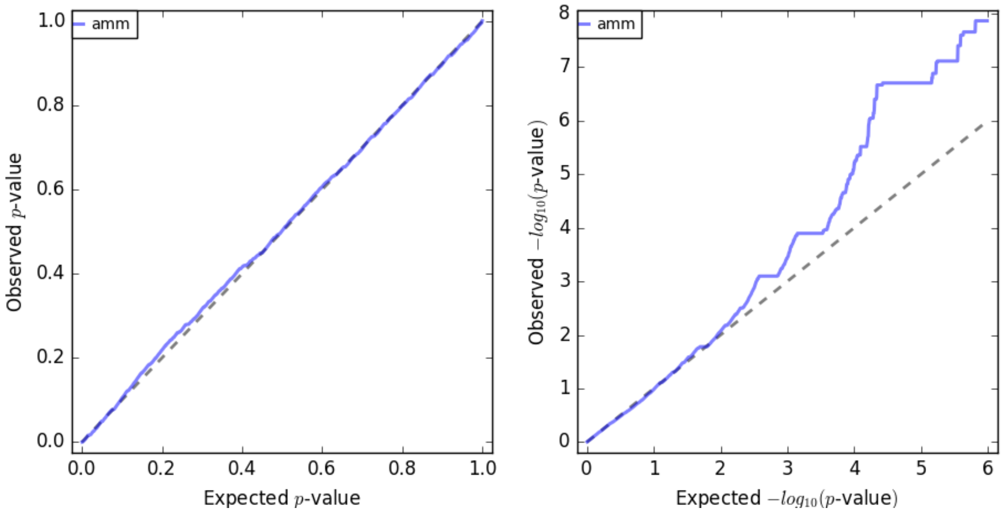

RAE

##### Linear model

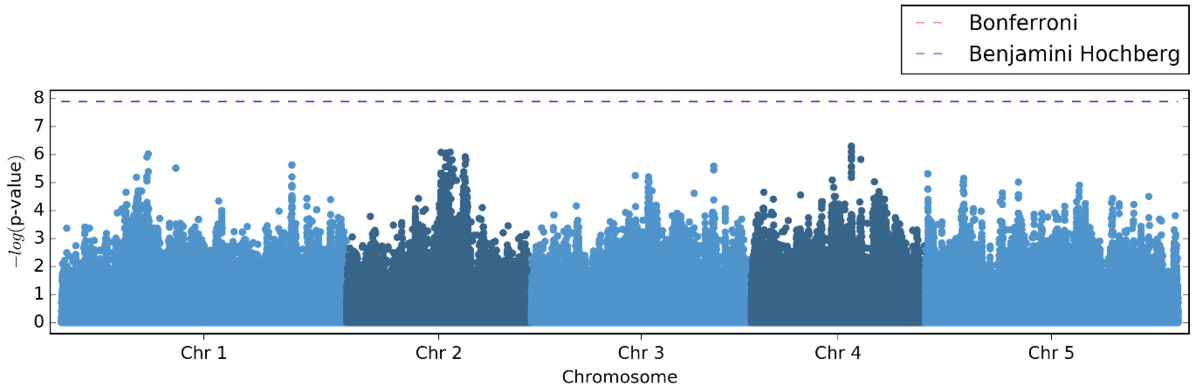


##### Kruskal-Wallis Model

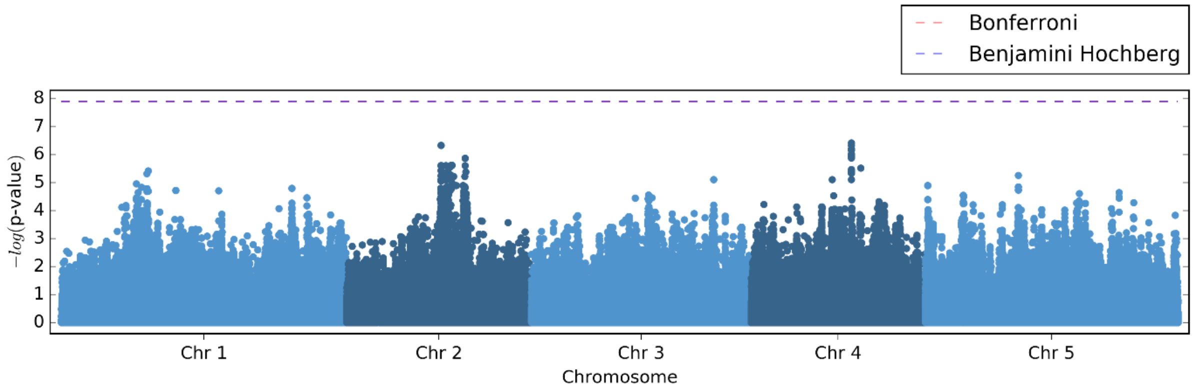


##### Accelerated Mixed Model

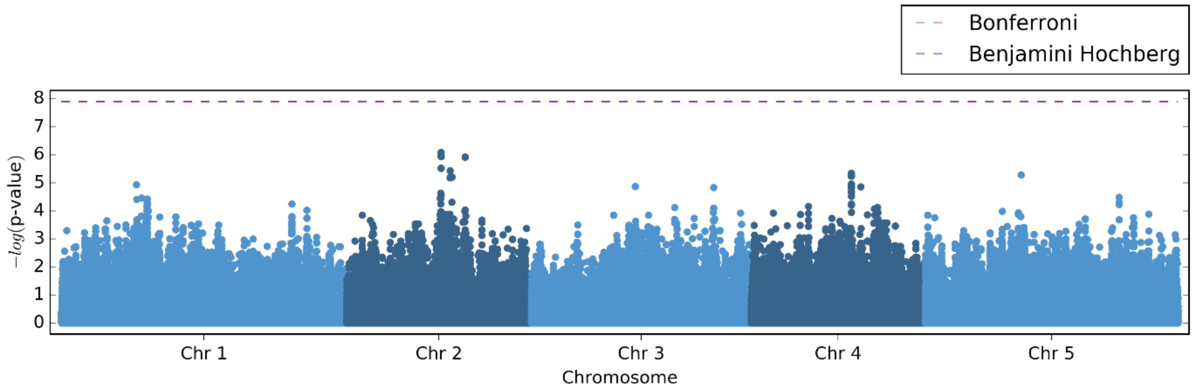

#### Linear model

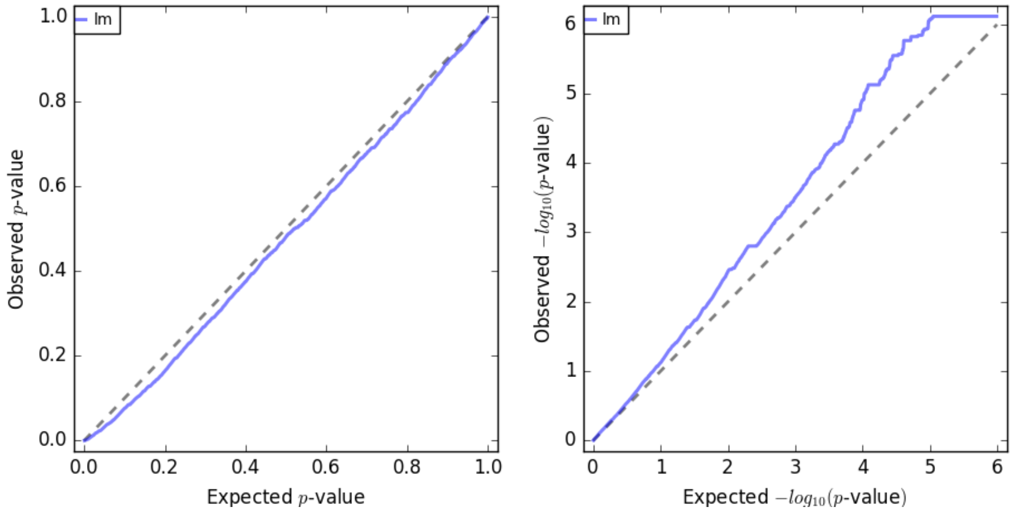


#### Kruskal-Wallis Model

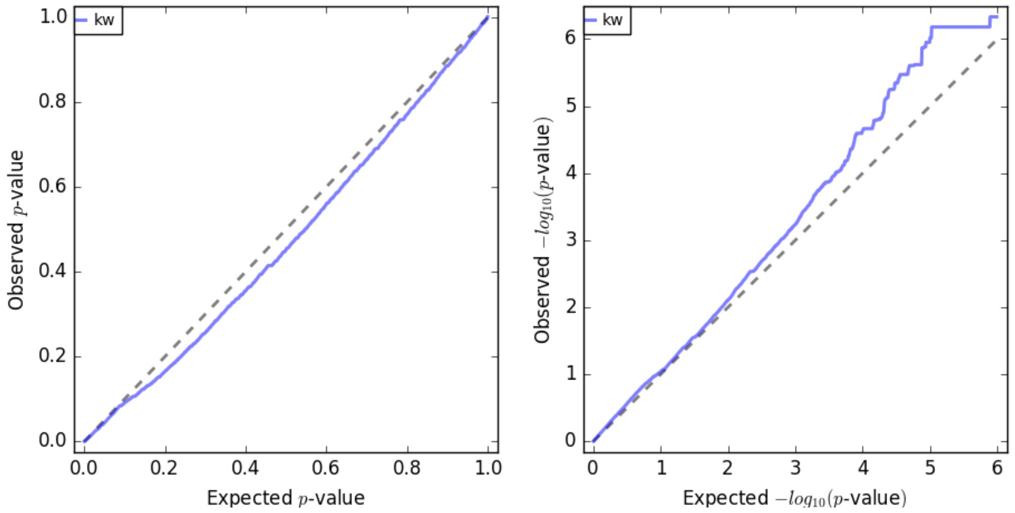


#### Accelerated Mixed Model

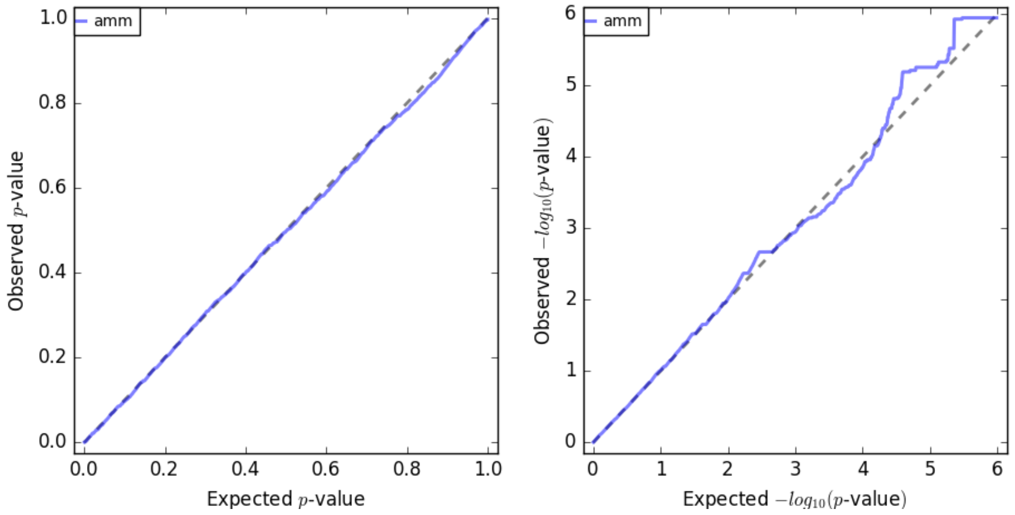

### Phase

#### Linear model

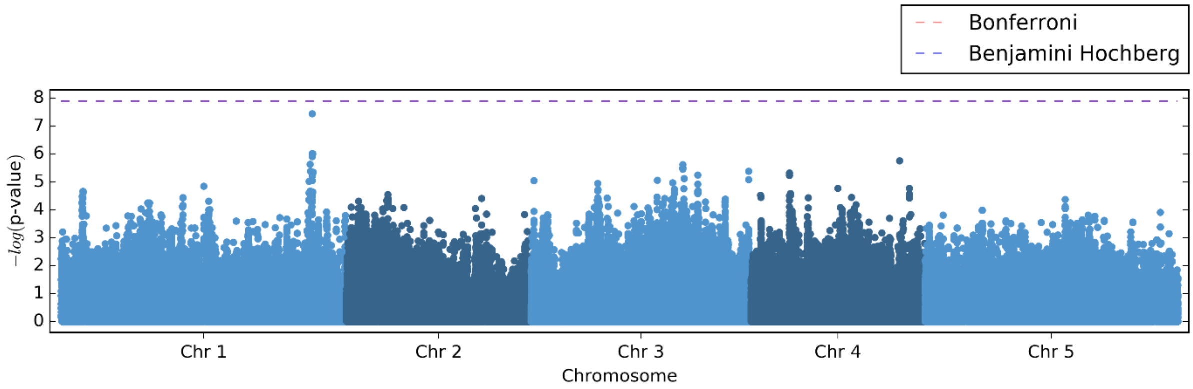


#### Kruskal-Wallis Model

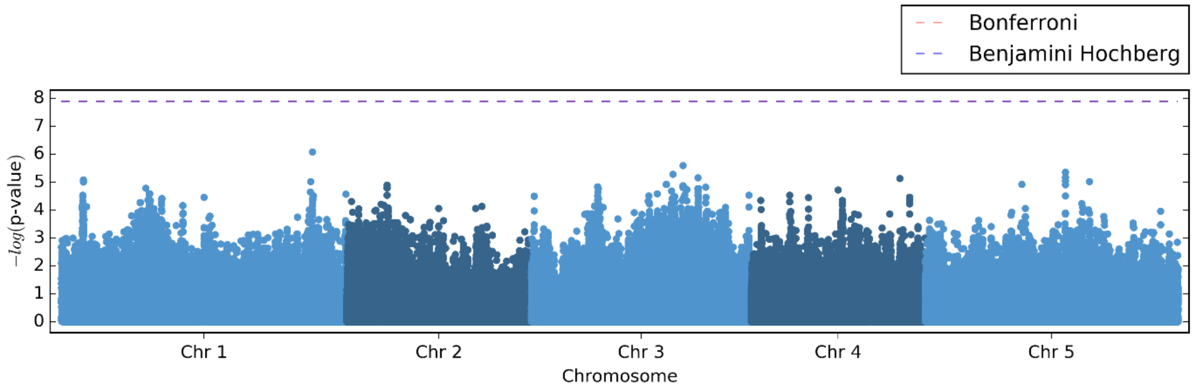


#### Accelerated Mixed Model

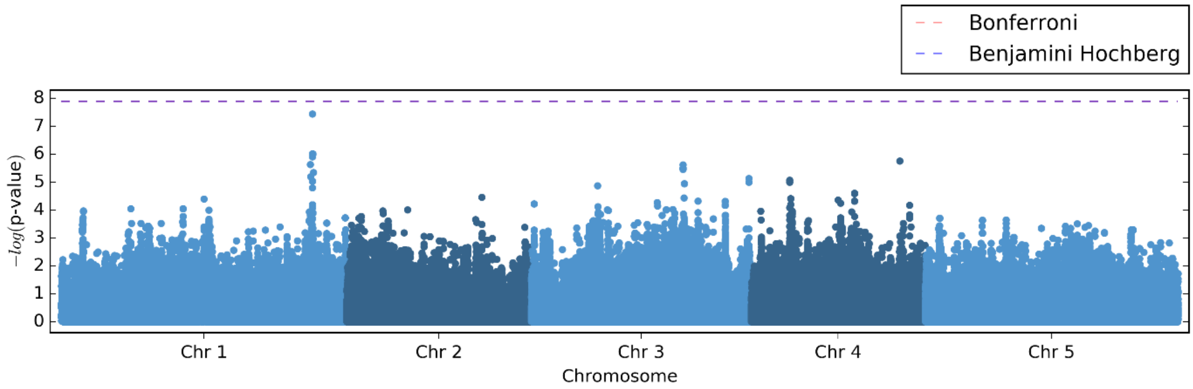

**Linear model**

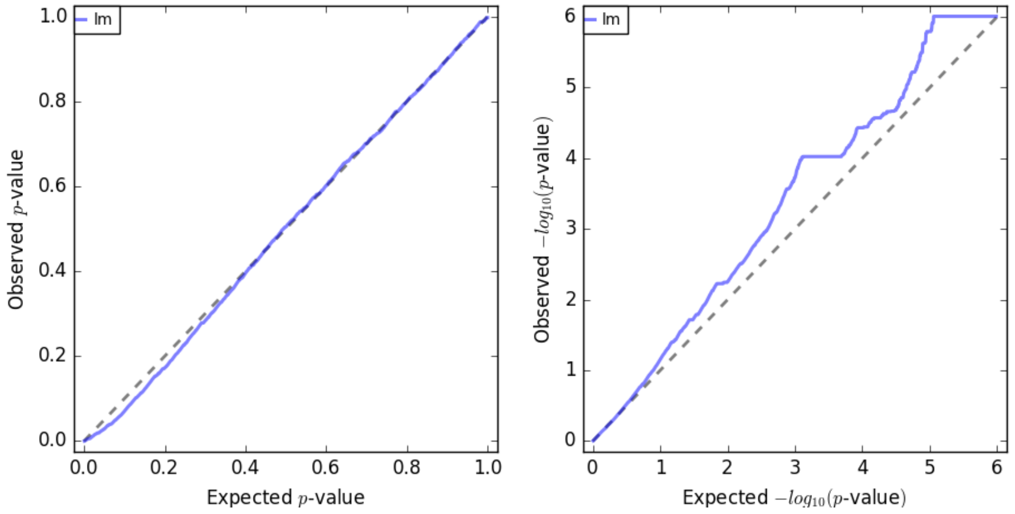


**Kruskal-Wallis Model**

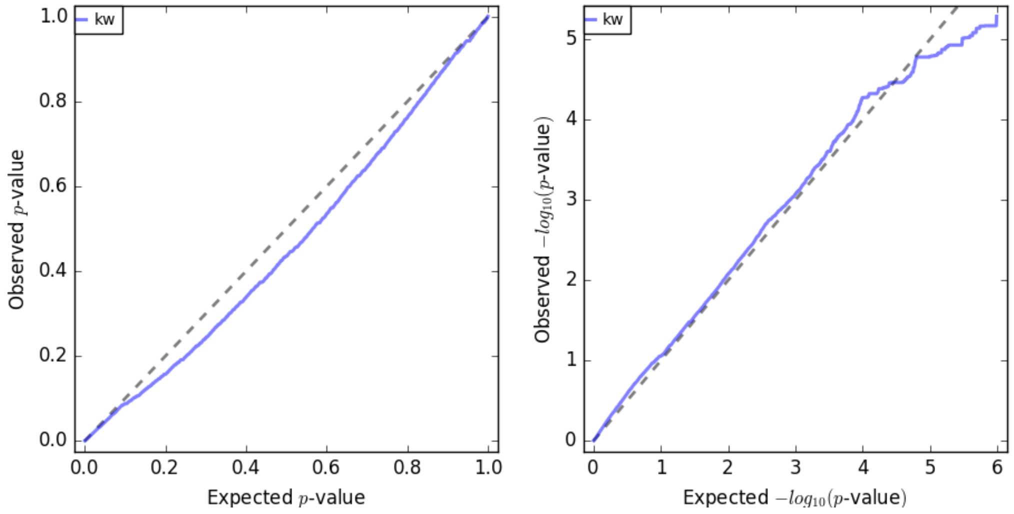


**Accelerated Mixed Model**

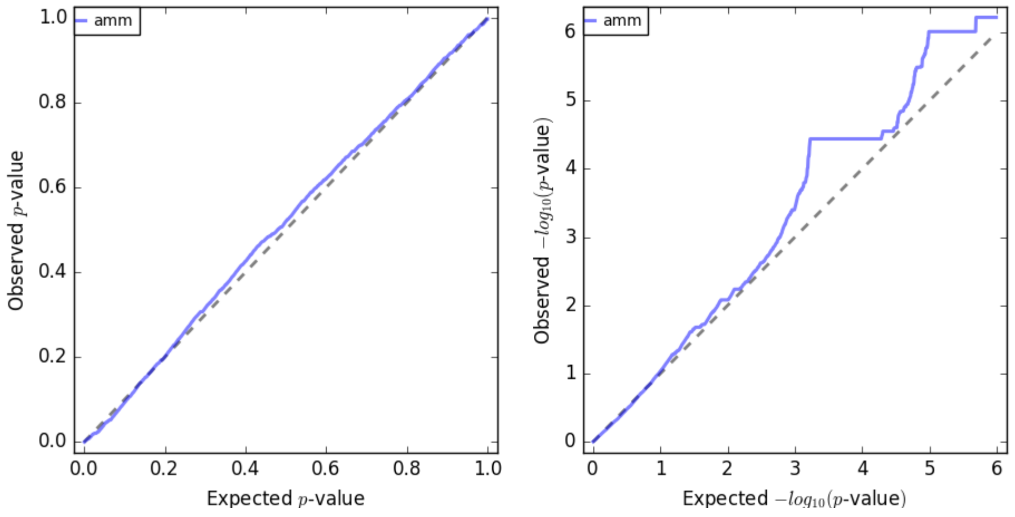

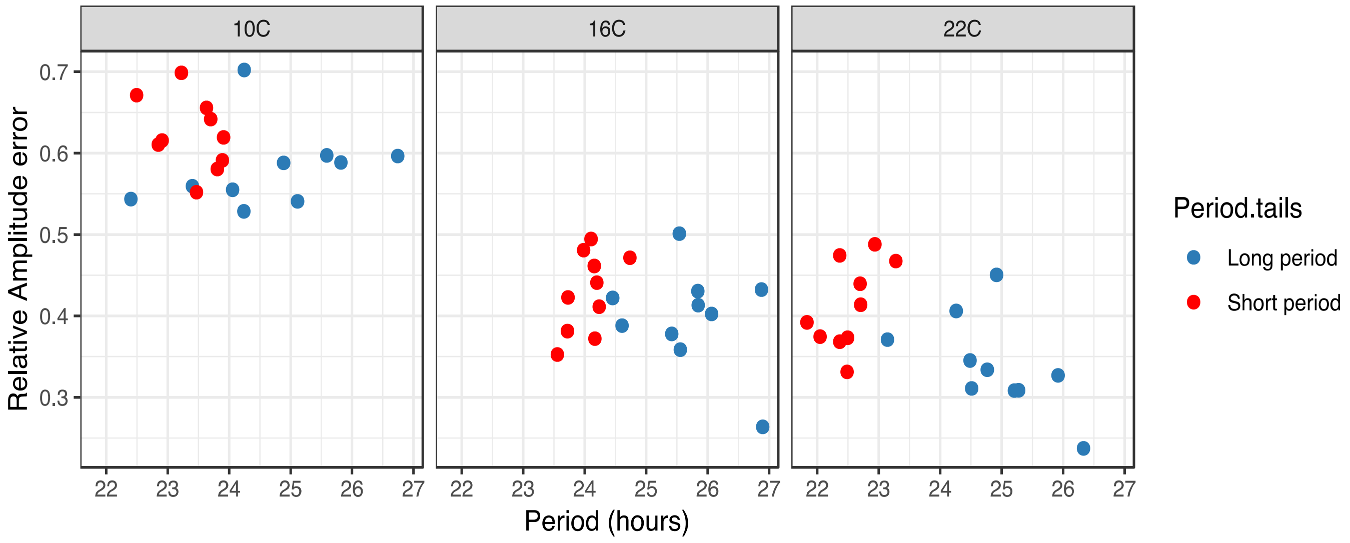


**Supplementary Figure 7. Period and RAE correlations in Period tail accessions**
Mean periods and RAE values were calculated using a general linear model for each accession in the period tails split across the three temperatures. The two tails remain split in period length across the three temperatures but rhythms are less robust at 10C (higher RAE).

A

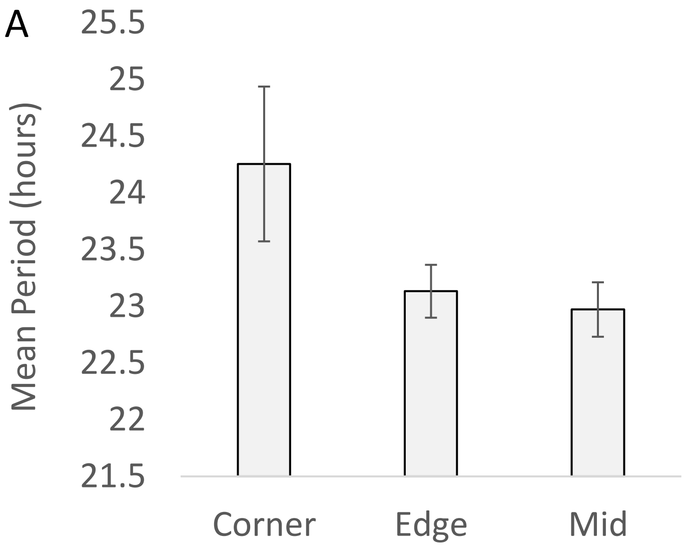


B

| | 1 | 2 | 3 | 4 | 5 | 6 | 7 | 8 | 9 | 10 | 11 | 12 |
|---|---|---|---|---|---|---|---|---|---|---|---|---|
| A | 23.98 | 22.95 | 23.36 | 22.95 | 23.67 | 23.62 | 23.36 | 23.29 | 23.25 | 22.81 | 23.42 | 25.24 |
| B | 23.52 | 23.05 | 23.37 | 22.78 | 22.86 | 23.26 | 23.16 | 22.91 | 22.61 | 23.03 | 22.67 | 23.26 |
| C | 23.28 | 22.98 | 23.16 | 22.73 | 22.92 | 22.76 | 23.29 | 22.45 | 23.26 | 23.12 | 23.16 | 23.05 |
| D | 22.94 | 23.06 | 22.80 | 22.54 | 22.86 | 22.90 | 22.99 | 23.00 | 23.22 | 22.89 | 23.58 | 23.05 |
| E | 23.08 | 23.29 | 22.85 | 23.12 | 22.94 | 22.72 | 22.99 | 22.95 | 22.94 | 23.25 | 22.92 | 23.13 |
| F | 23.10 | 23.08 | 22.59 | 22.68 | 22.82 | 23.10 | 23.15 | 22.96 | 23.24 | 22.97 | 23.19 | 23.25 |
| G | 22.93 | 22.83 | 23.18 | 22.79 | 23.16 | 22.55 | 22.83 | 23.53 | 22.83 | 22.64 | 22.85 | 23.12 |
| H | 24.12 | 22.96 | 22.81 | 23.27 | 22.85 | 22.96 | 23.04 | 22.73 | 23.08 | 22.96 | 23.19 | 23.68 |

**Supplementary Figure 8. Position in 96-well plate affects period estimation-justification for removing these wells from analysis.**

Mean periods for each well of all 96-well plates used in the 191 accession phenotyping dataset were calculated in Genstat. Wells were categorised into ‘corner’ ‘edge’ or ‘middle’ wells and group means were then plotted in the bar chart above (A). Error bars are standard deviation.

Corner wells have much longer period estimates than internal wells irrespective of accession used. Figure B shows all well-means colour-coded for mean period. (Red is long period, purple is short).

Several factors could be affecting period in these wells; they dry out fastest due to exposure to the external atmosphere and less contact with water condensation which could directly influence circadian rhythms. It is also possible that the corner wells are shielded from the overhead light by the side of the plate and so are receiving lower light levels than in the rest of the plate.

REML component effects: Period, RAE, Log10Amplitude

| 191 Accessions dataset- Y variate: Period | | | | | | |
|---|---|---|---|---|---|---|
| | | Wald statistic | n.d.f | F statistic | d.d.f | F pr |
| Fixed effects | Cabinet | 22.41 | 1 | 22.41 | 12.2 | <0.001 |
| | Run | 10.99 | 2 | 5.5 | 12.3 | 0.02 |
| | Cabinet.Run | 9.83 | 2 | 4.92 | 12.2 | 0.027 |
| Random effects | | Variance component | s.e. | | | |
| | Replicate | 0.022 | 0.015 | | | |
| | Replicate.block | 0.022 | 0.015 | | | |
| | Accession ID | 0.611 | 0.076 | | | |
| | residual | 1.539 | 0.046 | | | |

**Supplementary Table 1. Output from Genstat using the REML directive on 191 period data** Using a mixed linear model with cabinet and experimental run as fixed effects and replicate and Accession ID as random effects. n.d.f., numerator degrees of freedom (df); d.d.f., denominator degrees of freedom; F pr, F-test P-value.

<table>
<tr><td colspan="6">191 Accessions dataset- Y variate: RAE</td><td></td></tr>
<tr><td></td><td></td><td>Wald statistic</td><td>n.d.f</td><td>F statistic</td><td>d.d.f</td><td>F pr</td></tr>
<tr><td rowspan="3">Fixed effects</td><td>Cabinet</td><td>0.07</td><td>1</td><td>0.07</td><td>12.1</td><td>0.8</td></tr>
<tr><td>RUN</td><td>11.72</td><td>2</td><td>5.86</td><td>12.1</td><td>0.017</td></tr>
<tr><td>Cabinet.RUN</td><td>1.12</td><td>2</td><td>0.56</td><td>12.1</td><td>0.585</td></tr>
<tr><td rowspan="5">Random effects</td><td></td><td>Variance component</td><td>s.e.</td><td colspan="3" rowspan="5"></td></tr>
<tr><td>Replicate</td><td>0.00018</td><td>0.00013</td></tr>
<tr><td>Replicate.block</td><td>0.00039</td><td>0.00015</td></tr>
<tr><td>Accession ID</td><td>0.00114</td><td>0.00022</td></tr>
<tr><td>residual</td><td>0.0127</td><td>0.00038</td></tr>
</table>

**Supplementary Table 2. Output from Genstat using the REML directive on 191 RAE data**
Using a mixed linear model with cabinet and experimental run as fixed effects and replicate and Accession ID as random effects. n.d.f., numerator degrees of freedom (df); d.d.f., denominator degrees of freedom; F pr, F-test P-value.

| 191 Accessions dataset- Y variate: log10Amplitude | | | | | | |
|---|---|---|---|---|---|---|
| | | Wald statistic | n.d.f | F statistic | d.d.f | F pr |
| Fixed effects | Cabinet | 2.83 | 1 | 2.83 | 12 | 0.118 |
| | RUN | 377.1 | 2 | 188.55 | 12 | <0.001 |
| | Cabinet.RUN | 60.58 | 2 | 30.29 | 12 | <0.001 |
| Random effects | | Variance component | s.e. | | | |
| | Replicate | 0.0054 | 0.0038 | | | |
| | Replicate.block | 0.022 | 0.0038 | | | |
| | Accession ID | 0.0005 | 0.0012 | | | |
| | residual | 0.138 | 0.0042 | | | |

**Supplementary Table 3. Output from Genstat using the REML directive on 191 log10Amplitude data**

Using a mixed linear model with cabinet and experimental run as fixed effects and replicate and Accession ID as random effects. n.d.f., numerator degrees of freedom (df); d.d.f., denominator degrees of freedom; F pr, F-test P-value.

Likelihood testing: Period, Phase, RAE, Log10Amp

Accession omitted: Period~Cabinet+Run+Cabinet.Run + Replicate+ Replicate.block
Accession included: Period~Cabinet+Run+Cabinet.Run + Replicate+ Replicate.block + Accession ID

| PERIOD | | | | | | |
|---|---|---|---|---|---|---|
| | d.f. | AIC | Deviance | Chi Sq | Chi Df | Pr(Chisq) |
| omitted | 2516 | 4489.44 | 4483.44 | | | |
| included | 2515 | 4035.87 | 4027.87 | 455.57 | 1 | <0.0001 |

**Supplementary Table 4.** Likelihood ratio test and Akaike information criterion for effect of accession on period.
Comparison of two models for period with and without Accession ID included as a random effect. d.f., degrees of freedom; AIC, Akaike Information Criterion; Chi Sq, chi-squared statistic comparing the two models; Chi Df, difference in df of the two models; Pr(Chisq), chi-squared test P-value.

Accession omitted: Phase~Cabinet+Run
Accession included: Phase~Cabinet+Run+Accession ID

| PHASE | | | | | | |
|---|---|---|---|---|---|---|
| | d.f. | AIC | Deviance | Chi Sq | Chi Df | Pr(Chisq) |
| omitted | 2517 | -2419 | -2421 | | | |
| included | 2326 | -2824 | -2828 | 407 | 191 | <0.0001 |

**Supplementary Table 5.** Likelihood ratio test and Akaike information criterion for effect of accession on phase.
Comparison of two models for phase with and without Accession ID included as a random effect. d.f., degrees of freedom; AIC, Akaike Information Criterion; Chi Sq, chi-squared statistic comparing the two models; Chi Df, difference in df of the two models; Pr(Chisq), chi-squared test P-value.

Accession omitted: RAE~Cabinet+Run+Cabinet.Run + Replicate+ Replicate.block
Accession included: RAE~Cabinet+Run+Cabinet.Run + Replicate+ Replicate.block + Accession ID

| RAE | | | | | | |
|---|---|---|---|---|---|---|
| | d.f. | AIC | Deviance | Chi Sq | Chi Df | Pr(Chisq) |
| omitted | 2516 | -8156.68 | -8162.68 | | | |
| included | 2515 | -8220.65 | -8228.65 | 65.97 | 1 | <0.0001 |

**Supplementary Table 6.** Likelihood ratio test and Akaike information criterion for effect of accession on RAE.
Comparison of two models for RAE with and without Accession ID included as a random effect. d.f., degrees of freedom; AIC, Akaike Information Criterion; Chi Sq, chi-squared statistic comparing the two models; Chi Df, difference in df of the two models; Pr(Chisq), chi-squared test P-value.

Accession omitted: Log10Amp~Cabinet+Run+Cabinet.Run + Replicate+ Replicate.block
Accession included: Log10Amp ~Cabinet+Run+Cabinet.Run + Replicate+ Replicate.block + Accession ID

| LOG10(Amplitude) | | | | | | |
|---|---|---|---|---|---|---|
| | d.f. | AIC | Deviance | Chi Sq | Chi Df | Pr(Chisq) |
| omitted | 2516 | -2219.52 | -2225.52 | | | |
| included | 2515 | -2217.71 | -2225.71 | 0.19 | 1 | 0.663 |

**Supplementary Table 7.** Likelihood ratio test and Akaike information criterion for effect of accession on Log10Amplitude.
Comparison of two models for Log10Amplitude with and without Accession ID included as a random effect. d.f., degrees of freedom; AIC, Akaike Information Criterion; Chi Sq, chi-squared statistic comparing the two models; Chi Df, difference in df of the two models; Pr(Chisq), chi-squared test P-value.

**Supplementary Table 8. Accessions in Period Tails for temperature experiments**

| Accession ID | Accession name | Short or long tail | Accession Mean Period | Accession SE Period |
|---|---|---|---|---|
| 8387 | St-0 | Short | 21.278 | 0.329 |
| 6043 | Lšv-1 | Short | 21.459 | 0.329 |
| 6240 | TOM 06 | Short | 21.462 | 0.300 |
| 1552 | Sku-30 | Short | 21.568 | 0.370 |
| 6153 | TAA 03 | Short | 21.612 | 0.319 |
| 6030 | Gršn-5 | Short | 21.651 | 0.309 |
| 6201 | TDr-16 | Short | 21.723 | 0.341 |
| 9343 | Dju-1 | Short | 21.725 | 0.292 |
| 9433 | Nyl 13 | Short | 21.762 | 0.387 |
| 6238 | TOM 04 | Short | 21.837 | 0.341 |
| 9404 | HolA-1 1 | Long | 24.229 | 0.309 |
| 9453 | Stenk-2 | Long | 24.267 | 0.309 |
| 6022 | Fjä2-6 | Long | 24.331 | 0.329 |
| 6413 | Ull3-4 | Long | 24.357 | 0.292 |
| 6096 | T1060 | Long | 24.388 | 0.319 |
| 6035 | Hov1-10 | Long | 24.448 | 0.319 |
| 6114 | T570 | Long | 24.678 | 0.285 |
| 6149 | T970 | Long | 24.955 | 0.387 |
| 6123 | T680 | Long | 25.539 | 0.319 |
| 6133 | T800 | Long | 25.700 | 0.355 |

**Supplementary Table 9. Accessions in Phase Tails for temperature experiments**

| Accession ID | Accession name | Dusk or Dawn phase Tail | Accession Mean circular Phase | Accession SE Phase |
|---|---|---|---|---|
| 6011 | Eden-6 | Dusk | 12.21 | 0.58 |
| 6123 | T680 | Dusk | 13.93 | 0.23 |
| 6133 | T800 | Dusk | 14.07 | 0.23 |
| 6124 | T690 | Dusk | 14.48 | 0.21 |
| 6038 | Hov3-5 | Dusk | 15.01 | 0.24 |
| 9381 | Fri 1 | Dusk | 15.16 | 0.20 |
| 6035 | Hov1-10 | Dusk | 15.19 | 0.21 |
| 9332 | Bar 1 | Dusk | 15.27 | 0.21 |
| 6132 | T790 | Dusk | 15.30 | 0.27 |
| 6149 | T970 | Dusk | 15.72 | 0.22 |
| 6043 | Löv-1 | Dawn | 21.28 | 0.20 |
| 6013 | Eden-9 | Dawn | 21.34 | 0.29 |
| 6069 | Nyl-7 | Dawn | 21.45 | 0.19 |
| 8230 | Algutsrum | Dawn | 21.47 | 0.21 |
| 6074 | Ör-1 | Dawn | 21.51 | 0.20 |
| 9433 | Nyl 13 | Dawn | 21.74 | 0.23 |
| 1552 | Sku-30 | Dawn | 21.78 | 0.21 |
| 6036 | Hov3-2 | Dawn | 21.85 | 0.21 |
| 8387 | St-0 | Dawn | 21.90 | 0.20 |
| 9402 | Hel-3 | Dawn | 22.54 | 0.20 |

**Supplementary Table 10. Accessions in RAE Tails for temperature experiments**

| Accession ID | Accession name | RAE: low and high | Accession Mean RAE | Accession SE RAE |
|---|---|---|---|---|
| 6123 | T680 | Low | 0.310999 | 0.023396 |
| 6114 | T570 | Low | 0.316548 | 0.021708 |
| 1063 | Brösarp-21-140 | Low | 0.326776 | 0.021708 |
| 5832 | App1-16 | Low | 0.338294 | 0.021707 |
| 9404 | HolA-1 1 | Low | 0.343043 | 0.022936 |
| 6126 | T720 | Low | 0.34418 | 0.023396 |
| 6218 | TFÄ 08 | Low | 0.344506 | 0.021708 |
| 6173 | TÅD 05 | Low | 0.344854 | 0.022936 |
| 9383 | Fri 3 | Low | 0.346338 | 0.021708 |
| 6095 | T1050 | Low | 0.347766 | 0.022503 |
| 6210 | TEDEN 03 | High | 0.426781 | 0.025563 |
| 6238 | TOM 04 | High | 0.428421 | 0.024406 |
| 6010 | Eden-5 | High | 0.42886 | 0.02621 |
| 8387 | St-0 | High | 0.430485 | 0.023885 |
| 6043 | Löv-1 | High | 0.438462 | 0.023884 |
| 9436 | Puk-1 | High | 0.438879 | 0.022935 |
| 992 | Ale-Stenar-44-4 | High | 0.439084 | 0.025561 |
| 6070 | Omn-1 | High | 0.441196 | 0.026206 |
| 6209 | TEDEN 02 | High | 0.441694 | 0.024406 |
| 9402 | Hel-3 | High | 0.443792 | 0.023883 |

| **Trait** | **Freezing tolerance** | | **Flowering time at 10C** | | **Flowering time at 16C** | | **Flowering time (Mean in Swedish environment)** | | **Flowering time (Mean in Spanish environment)** | | **Seed dormancy** | |
|---|---|---|---|---|---|---|---|---|---|---|---|---|
| Source | Horton 2016 | | Sasaki 2015 | | Sasaki 2015 | | Li 2010 | | Li 2010 | | Kerdaffrec 2016 | |
| N in common | 107 | | 127 | | 127 | | 29 | | 29 | | 123 | |
| Circadian traits | $R^2$ | *p* | $R^2$ | *p* | $R^2$ | *p* | $R^2$ | *p* | $R^2$ | *p* | $R^2$ | *p* |
| Period | 0.008 | 0.352 | 0.027 | 0.063 | 0.001 | 0.708 | 0.227 | **0.009** | 0.163 | 0.030 | 0.008 | 0.352 |
| Phase | 0.010 | 0.317 | 0.045 | 0.017 | 0.009 | 0.299 | 0.256 | **0.005** | 0.241 | **0.007** | 0.010 | 0.317 |
| RAE | 0.006 | 0.446 | 0.005 | 0.417 | 0.000 | 0.838 | 0.058 | 0.208 | 0.020 | 0.461 | 0.006 | 0.446 |

**Supplementary Table 11. Linear regression of circadian data from 191 Swedish Arabidopsis accessions with common accessions from previously published datasets**. Significant *p*-values (<0.01) are highlighted in red. Data from Sasaki et al was measured under long days (16h-8h) and constant temperatures in controlled growth cabinets. Data from Li et al is the mean of four conditions replicating either Swedish or Spanish weather patterns across two years.

Mutant validation tables

**Supplementary Table 12:** Testing differences in period between candidate circadian mutants and their respective controls (highlighted in grey) (Welch Two Sample t-test)

| Plant ID | Sample size across all reps | Period mean (hours) | Period SD | t value | df | p value |
|---|---|---|---|---|---|---|
| **col-0** | 180 | **24** | **2.02** | | | |
| *cor27-1* | 108 | 24 | 1.75 | 0.18 | 118 | 0.857 |
| *cor27-2* | 107 | 24.2 | 2.21 | -0.35 | 142 | 0.726 |
| *cor28-2* | 158 | 25.3 | 1.78 | -3.86 | 117 | >0.001 |
| *cor28-2/27-1* | 101 | 26.5 | 2.33 | -6.24 | 139 | >0.001 |
| *mybl2* | 103 | 24 | 1.97 | -0.06 | 140 | 0.955 |
| *parc6* | 97 | 23.9 | 2.08 | 0.36 | 138 | 0.722 |
| *parc6-1* | 104 | 23.4 | 1.77 | 1.93 | 120 | 0.056 |
| *atg1g71015* | 102 | 23.7 | 2.11 | 0.96 | 138 | 0.338 |
| **WT for cor28-1** | 72 | **23.9** | **1.62** | | | |
| *cor28-1* | 70 | 25.3 | 1.73 | 2.41 | 39 | 0.021 |
| **ler** | 108 | **23.2** | **2.55** | | | |
| *sco2* | 101 | 23.7 | 1.95 | -0.61 | 92 | 0.546 |

**Supplementary Table 13:** Testing differences in RAE between candidate circadian mutants and their respective controls (highlighted in grey) (Welch Two Sample t-test)

| Plant ID | Sample size across all reps | RAE mean | RAE SD | t value | df | p-value |
|---|---|---|---|---|---|---|
| **col-0** | 180 | **0.42** | **0.18** | | | |
| *cor27-1* | 108 | 0.34 | 0.18 | 3.7612 | 132.65 | >0.001 |
| *cor27-2* | 107 | 0.29 | 0.14 | 6.1329 | 109.28 | >0.001 |
| *cor28-2* | 158 | 0.28 | 0.13 | 6.2523 | 104.52 | >0.001 |
| *cor28-2/27-1* | 101 | 0.28 | 0.14 | 5.7682 | 120.7 | >0.001 |
| *mybl2* | 103 | 0.37 | 0.17 | 2.6062 | 128.76 | 0.01024 |
| *parc6* | 97 | 0.37 | 0.16 | 3.0367 | 126.56 | 0.00291 |
| *parc6-1* | 104 | 0.35 | 0.17 | 3.6523 | 130.41 | >0.001 |
| *atg1g71015* | 102 | 0.38 | 0.16 | 2.6593 | 124.82 | >0.001 |
| **WT for cor28-1** | 72 | **0.33** | **0.17** | | | |
| *cor28-1* | 70 | 0.28 | 0.16 | -0.87301 | 36.889 | 0.3883 |
| **ler** | 108 | **0.43** | **0.17** | | | |
| *sco2* | 101 | 0.51 | 0.16 | -1.7234 | 120.37 | 0.08738 |

**Supplementary Table 14:** Testing differences in Phase between candidate circadian mutants and their respective controls (highlighted in grey) (Watson's Two-Sample Test of Homogeneity)

| *Plant ID* | N | Circular Phase mean (hours) | Phase SD | test statistic | P value |
|---|---|---|---|---|---|
| **col-0** | 180 | 15.35 | 1.04 | | |
| *cor27-1* | 108 | 13.39 | 0.77 | 0.4726 | < 0.001 |
| *cor27-2* | 107 | 14.02 | 0.83 | 0.2467 | < 0.05 |
| *cor28-2* | 158 | 12.93 | 0.91 | 0.3669 | < 0.01 |
| *cor28-2/27-1* | 101 | 12.67 | 0.98 | 0.6332 | < 0.001 |
| *mybl2* | 103 | 14.27 | 0.93 | 0.0911 | > 0.10 |
| *parc6* | 97 | 13.91 | 0.97 | 0.0886 | > 0.10 |
| *parc6-1* | 104 | 14.95 | 0.87 | 0.0599 | > 0.10 |
| *atg1g71015* | 102 | 14.45 | 0.92 | 0.1455 | > 0.10 |
| **WT for cor28-1** | 72 | 14.54 | 0.79 | | |
| *cor28-1* | 70 | 12.95 | 0.86 | 0.1941 | < 0.05 |
| **ler** | 108 | 16.37 | 1.04 | | |
| *sco2* | 101 | 15.33 | 1 | 0.0568 | > 0.10 |

**Supplementary Table 15. Accumulated analysis of variance table for period with temperature data**

| **Period** | | | | |
|---|---|---|---|---|
| **(Cabinet+Temp)*Period_tails/accession ID** | | | | |
| Accumulated analysis of variance | | | | |
| Change | df | Mean Square | Variance ratio | F probablility |
| Cabinet | 1 | 52.621 | 10.71 | 0.001 |
| Temp | 2 | 165.812 | 33.74 | <.001 |
| Period_tails | 1 | 1008.214 | 205.15 | <.001 |
| Cabinet*Period_tails | 1 | 49.504 | 10.07 | 0.002 |
| Temp*Period_tails | 2 | 35.499 | 7.22 | <.001 |
| Period_tails* accession ID | 18 | 23.015 | 4.68 | <.001 |
| Cabinet*Period_tails* accession ID | 18 | 3.921 | 0.80 | 0.705 |
| Temp*Period_tails* accession ID | 36 | 6.697 | 1.36 | 0.076 |
| Residual | 1146 | 4.915 | | |
| Total | 1225 | 6.425 | | |

**Supplementary Table 16. Accumulated analysis of variance table for RAE with temperature data**

| **RAE** | | | | |
|---|---|---|---|---|
| **(Cabinet+Temp)*RAE_tails/Accession ID** | | | | |
| Accumulated analysis of variance | | | | |
| Change | df | Mean Square | Variance ratio | F probablility |
| Cabinet | 1 | 0.2799 | 9.52 | 0.002 |
| Temp | 2 | 5.60176 | 190.48 | <.001 |
| RAE_tails | 1 | 2.08938 | 71.05 | <.001 |
| Cabinet*RAE_tails | 1 | 0.08555 | 2.91 | 0.088 |
| Temp*RAE_tails | 2 | 0.34449 | 11.71 | <.001 |
| RAE_tails* accession ID | 18 | 0.12326 | 4.19 | <.001 |
| Cabinet*RAE_tails* accession ID | 18 | 0.02225 | 0.76 | 0.753 |
| Temp*RAE_tails* accession ID | 36 | 0.03589 | 1.22 | 0.176 |
| Residual | 1148 | 0.02941 | | |
| Total | 1227 | 0.0424 | | |

**Supplementary Table 17. Circular regression analysis for Phase with temperature data**

| **Phase** | | | | | | |
|---|---|---|---|---|---|---|
| **Circular regression analysis** | | | | | | |
| Response variate: Degrees | | | | | | |
| Distribution: von Mises | | | | | | |
| Link function: Mu = Mu0 + 2*ARCTAN(liniar prediction) | | | | | | |
| Number of units: 1107 | | | | | | |
| **Fitted terms: Cabinet + Temp + Phase_tails + Temp.Phase_tails +Phase_tails. accession ID + Temp.Phase_tails.accession ID** | | | | | | |
| circular regression model | removed term | d.f | Deviance | Diffierence in Deviance | difference in d.f | P-value |
| Full model:Cabinet + Temp + Phase_tails + Temp.Phase_tails +Phase_tails. accession ID + Temp.Phase_tails. accession ID | | 55 | -790.4 | | | |
| Cabinet + Temp + Phase_tails + Temp.Phase_tails +Phase_tails. accession ID | Temp.Phase _tails. **accession ID** | 23 | -725.2 | 65.2 | 32 | <0.001 |
| Cabinet + Temp + Phase_tails + Temp.Phase_tails | Phase_tails. **accession ID** | 7 | -691.0 | 34.2 | 16 | <0.01 |
| Cabinet + Temp + Phase_tails | Temp.Phase _tails | 5 | -689.7 | 1.3 | 2 | 0.52 |
| Cabinet + Temp | Phase_tails | 4 | -656.4 | 33.3 | 1 | <0.00001 |
| Temp | Cabinet | 3 | -653.7 | 2.7 | 1 | 0.1 |
| Null | Temp | | | 231.7 | 3 | <0.00001 |
