## Supplementary File 3. Line means and SD for temperature experiments for "Circadian diversity in Swedish *Arabidopsis* accessions is associated with naturally occurring genetic variation in *COR28*"

Selecting ROI for DF imaging using multiple 96 well plates

1. Open stack of images using FIJI. Transform stack to the correct orientation, add a fire Lookup table effect. HINT: This procedure works best on perfectly straight images. If horizontal lines in your image are not aligned with the window edges, use the segmented line tool to create a reference line and then use Edit>Selection>Straighten with line width at approx. 600 pixels.
2. Import the ‘circle tool’ macro. Right click on macro icon and select radius size (approx. 10)

*// This macro set demonstrates how a tool*

*// can configures by double clicking on it.*

*var radius = 20;*

*macro "Circle Tool - C00cO11cc" {*

*getCursorLoc(x, y, z, flags);*

*makeOval(x-radius, y-radius, radius*2, radius*2);*

*}*

*macro "Circle Tool Options" {*

*radius = getNumber("Radius: ", radius);*

*}*

3. Make sure stack image is sufficiently contrasted to be able to see each well. Using the circle tool, add a ROI to each top left hand corner of each plate.
4. Run the macro ‘generate_all_rois_corner_selection.ijm’. (Drag and drop text file onto FIJI toolbar.)

*getSelectionBounds(x, y, width, height)*

*print(x,y,width,height)*

*first_x = x*

*first_y = y*

*radius_in = width*

*radius = radius_in*

*col_gap = radius_in+0.5*

*row_gap = radius_in*

*start_x = first_x*

*start_y = first_y*

*for (row=0; row < 8; row++) {*

*for (col=0; col < 12; col++) {*

*makeOval(start_x + col * col_gap, start_y + row * row_gap, radius, radius);*

*roiManager("add");*

*}*

*}*

5. This will give you all of the ROIs but they might not be exactly where they should. For plates which need adjusting, select the original plate corner in the ROI manager then click the stack window and use arrow keys to nudge the ROI until it is over the wells.
6. Repeat with all corners which need re-doing. Delete all other ROIs and run the macro again.
7. Individual wells can be nudged independently of the whole plate by selecting in the manager and using the translate tool.
8. Save the ROI when happy with all positions. Save the transformed stack. Select all ROIs apart from the starting corners and multi measure to get integrated densities for each well. (ID must be ticked in Set measurements).
